# Local traveling waves of cytosolic calcium elicited by defense signals or wounding are propagated by distinct mechanisms

**DOI:** 10.1101/2025.01.23.634554

**Authors:** Weiwei Zhang, Nilay Kumar, Jessica R. Helwig, Alexis Hoerter, Anjali S. Iyer-Pascuzzi, David M. Umulis, Elsje Pienaar, Christopher J. Staiger

**Affiliations:** Department of Botany and Plant Pathology, Purdue University, West Lafayette, IN, United States; Weldon School of Biomedical Engineering, Purdue University, West Lafayette, IN, United States; Department of Biological Sciences, Purdue University, West Lafayette, IN, United States; EMBRIO Institute, Purdue University, West Lafayette, IN, United States

## Abstract

Cytosolic Ca^2+^ signatures with specific spatiotemporal patterns play crucial roles in plant responses to biotic and abiotic stresses. Perception of microbe- or damage-associated molecular patterns (MAMPs or DAMPs) initiates signaling cascades that represent the first layer of plant defense against pathogens known as pattern-triggered immunity (PTI). During PTI, MAMP/DAMP-induced cytosolic Ca^2+^ fluxes serve as essential messengers in the initiation and transmission of defense signals at the cellular, whole organ, and systemic levels. However, the specific patterns of these Ca^2+^ signatures in response to different pathogen cues, and the mechanisms that encode them, remain largely unexplored. In this study, we quantitatively assessed Ca^2+^ signatures at the single-cell level as well as the local traveling Ca^2+^ waves induced by global treatment of Arabidopsis cotyledons with MAMPs or DAMPs. We demonstrated that MAMPs induced distinct local spatiotemporal Ca^2+^ responses in epidermal pavement cells, with Ca^2+^ traveling waves consistently initiated from a subset of cells and spreading in an approximately radial pattern. These local traveling waves propagated at a slow but constant speed of approximately 1 µm/s and spread to a limited number of neighboring cells. In contrast, wound-induced traveling waves displayed a diffusion-like decay pattern that moved rapidly away from the wounded cell but with diminishing speed over time and distance. Mathematical modeling supported a calcium-induced calcium release mechanism that could recapitulate the constant wave speed induced by MAMPs. These findings contribute to a deeper understanding of plant defense-related Ca^2+^ signaling mechanisms as well as how defense responses are spatially restricted within tissues.

**One Sentence Summary:** A slow, local traveling wave of calcium is initiated from a subset of epidermal cells during plant immune signaling.

## INTRODUCTION

Cytosolic calcium ions ([Ca^2+^]_cyt_) are among the most versatile intracellular second messengers in eukaryotes, playing a crucial role in signal transduction cascades as well as signal propagation between cells in multicellular organisms (*1, 2*). Fluctuations in [Ca^2+^]_cyt_ can take the form of spikes or periodic oscillations whose particular features of amplitude, duration and frequency often display highly specific spatiotemporal patterns that are referred to as “signatures” (*3, 4*). These [Ca^2+^]_cyt_ signatures are typically unique to the input stimulus as well as to the cell or tissue type perceiving the stimulus. Unique [Ca^2+^]_cyt_ signatures are encoded by the coordinated activities of channels and pumps that mediate Ca^2+^ entry into and removal from the cytosol in a precise spatiotemporal manner (*5*). Calcium signatures are then decoded by specific sets of Ca^2+^-binding proteins, which serve as Ca^2+^ sensors or effectors, to alter numerous downstream activities and cell functions in response to intrinsic and extrinsic cues (*6, 7*). Although numerous molecular components involved in Ca^2+^ signature encoding and decoding have been identified, how cells coordinate these events to achieve spatiotemporal specificity in stimulus-induced Ca^2+^ signatures, from single-cell responses to signal propagation at the multicellular level, remain largely unexplored for many key responses to biotic or abiotic stresses.

Plants, as sessile organisms, rely on robust signaling mechanisms to detect external stresses and initiate rapid defense or mitigation responses, from local to whole-plant levels. It is well known that diverse external stimuli, such as mechanical wounding, high light, or osmotic stress, trigger fast-moving Ca^2+^ waves that propagate over long distances to communicate with distal regions of the plant (*7–9*). These traveling waves play a critical role in systemic signaling, thereby enabling the plant to better prepare and adapt to various stresses. Systemic Ca^2+^ waves predominantly travel through the plant vascular tissue and can reach speeds up to several hundred µm/s, whereas local waves propagating at the tissue level, for example across epidermal pavement cells, are reported to be 10 to 100 times slower (*10–12*). The speed of Ca^2+^ waves has been suggested as another factor linked to signal specificity, potentially generating spatially- and temporally-distinct patterns of downstream responses at the tissue or organ level (*4*). A recent study demonstrated that mechano-stimulation of a single epidermal pavement cell in Arabidopsis, whether by touch or by release, induced two distinct types of local Ca^2+^ waves that traveled at different speeds and distances (*13*).

Several factors that contribute to systemic transmission of Ca^2+^ traveling waves in plants, including Ca^2+^-permeable ion channels, reactive oxygen species (ROS), apoplastic diffusion of amino acids that activate glutamate receptor-like (GLR) calcium channels, and the permeability of plasmodesmata (PD) have been identified (*8, 14–16*). However, whether and how these factors contribute to distinct Ca^2+^ signatures at the single-cell level or their role in local traveling wave patterns remain largely unknown.

Calcium signaling plays an essential role during the plant defense response against microbial pathogens (*17*). The first layer of plant defense, termed pattern-triggered immunity (PTI), is induced by recognition of microbe-, pathogen-, or damage-associated molecular patterns (MAMPs, PAMPs or DAMPs) through cell surface-localized pattern recognition receptors (PRRs) (*18, 19*). During PTI, an influx of [Ca^2+^]_cyt_ is among the earliest responses to initiate and transmit defense signals, occurs within minutes of signal perception, and activates numerous downstream responses including an apoplastic ROS burst, cytoskeletal remodeling, altered gene expression, stomatal closure, callose deposition, and production of phytohormones (*20–22*). Distinct [Ca^2+^]_cyt_ signatures with spatially- and temporally-specific patterns are thought to be induced in plant cells following the recognition of specific types of MAMPs or DAMPs and likely represent a mechanism that plants utilize to fine-tune the location, magnitude, and duration of immune responses (*17, 23, 24*). However, many previous studies measured cytosolic Ca^2+^ transients at the whole organ or seedling level and lacked the resolution to address signal specificity across different temporal and spatial scales. Recent advances with genetically-encoded Ca^2+^ indicators and fluorescence microscope imaging modalities enable the characterization of stimulus-induced [Ca^2+^]_cyt_ transients at high spatiotemporal resolution (*25*). Live-cell imaging with these Ca^2+^ sensors revealed that MAMP-induced Ca^2+^ transients exhibit oscillatory patterns in individual Arabidopsis epidermal pavement cells or stomatal guard cells (*26–28*). Nevertheless, [Ca^2+^]_cyt_ signature specificity, both at the single cell and tissue level with respect to the sites of MAMP perception as well as transmission to neighboring cells during PTI, remain largely undefined.

To address these knowledge gaps, we developed a high spatiotemporal resolution quantitative imaging pipeline and characterized single-cell Ca^2+^ signatures as well as the spatial spread of Ca^2+^ traveling waves induced by various MAMPs or DAMPs in epidermal pavement cells from detached Arabidopsis cotyledons. We demonstrated that PTI-induced Ca^2+^ signals originated from a limited number of initiation sites in the epidermal tissue, showed a distinct pattern of transients at the single-cell level, and that local traveling waves of Ca^2+^ propagated at a low but constant speed and dissipated after spreading to a limited number of neighboring cells. The constant speed of MAMP-induced local traveling waves was distinct from wound-induced traveling waves, which have been shown to propagate by simple diffusion. The potential mechanisms for the local traveling waves of Ca^2+^ induced by MAMPs were tested with mathematical modeling and support a calcium-induced calcium release (CICR) mechanism for wave transmission. Our findings will help unravel the underlying mechanisms of Ca^2+^ signal encoding and traveling wave propagation at the cell and tissue level, as well as how various types of stimulus-induced signaling specificity are achieved in plants.

## RESULTS

### MAMPs induce defined spatiotemporal Ca^2+^ responses in epidermal tissue

To measure the intracellular Ca^2+^ transients and signatures elicited by global MAMP treatment as well as signal propagation from cell to cell, we used the genetically-encoded, intensiometric R-GECO1 reporter (*27*), which demonstrates high sensitivity and allows imaging of stimulus-induced cytosolic calcium ([Ca^2+^]_cyt_) fluxes at single-cell resolution in plants. The Arabidopsis cotyledon represents a widely used experimental system for high resolution spatiotemporal imaging as well as plant defense-related cellular dynamics and signaling studies (*29–31*). Epidermal pavement cells on the abaxial side of detached 7-day-old cotyledons were imaged in a custom-made chamber based on the design described in Keinath et al. (*27*) (fig. S1). The chamber allowed rapid perfusion and global or uniform application of elicitors or treatments as well as bottom imaging with an inverted objective on a spinning-disk confocal microscope. Samples were imaged for at least 10 min at 5-s intervals, prior to addition of MAMP elicitors, to eliminate any laser illumination-induced Ca^2+^ responses. After applying the immunogenic peptide flg22 (*32*), we consistently detected [Ca^2+^]_cyt_ transients and intercellular Ca^2+^ waves in cotyledon epidermal cells within 2–5 minutes (Figs. 1 and 2). We also observed that Ca^2+^ signatures at the single-cell level exhibited an oscillatory pattern, with 4–5 peaks lasting for 15–20 min, when 1 µM flg22 was applied (Figs. 1 and 2). This response time and oscillatory pattern were similar to previous reports with various imaging settings or Ca^2+^ reporters (*24, 26–28*), thus validating our experimental set-up for high spatiotemporal resolution imaging of Ca^2+^ transients during the plant immune response. In addition, we demonstrated that the fluorescence changes detected were Ca^2+^ dependent, because pre-treatment with 500 µM LaCl_3_, a Ca^2+^ channel blocker, or with 25 µM BAPTA-AM, a cell-permeant Ca^2+^ chelator, markedly reduced the flg22-induced signal increases in epidermal cells (fig. S2). Moreover, no Ca^2+^ elevations were detected when a null mutant for the pattern-recognition receptor FLAGELLIN-SENSING2 (FLS2; *33*), *fls2* expressing R-GECO1, was imaged, demonstrating that the observed Ca^2+^ dynamic changes were specific to flg22 elicitation and recognition by its cognate receptor (fig. S3).

**Fig. 1.**
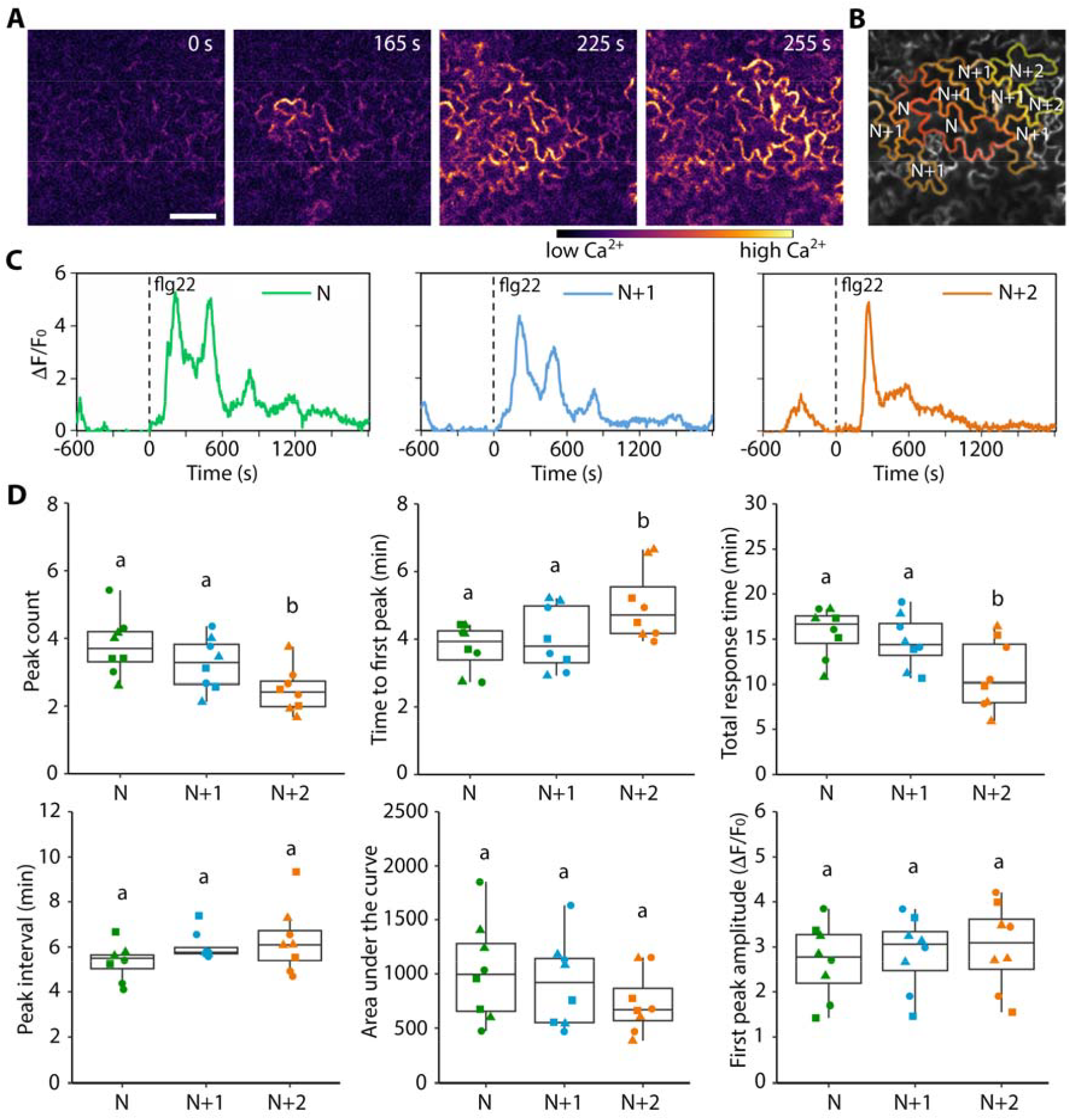
MAMP-induced Ca^2+^ signatures and cell-to-cell signal propagation follow a defined spatial pattern. (**A**) A representative timelapse image series of R-GECO1 fluorescence dynamics in Arabidopsis cotyledon epidermal cells following global treatment with 1 µM flg22. See also Supplemental Movie S1. Cytosolic Ca^2+^ elevations were first detected in initiator cells and then propagated to neighboring cells. Bar = 50 µm. (**B**) Annotation of initiator cells (N), N+1 and N+2 neighboring cells shown in (A). (**C**) Representative traces for the Ca^2+^ intensity ratio of an N, N+1, and N+2 cell, respectively. Additional representative traces are shown in Figure S5. Fluorescence intensity changes were calculated as the ratio ΔF/F_0_. Black dashed lines indicate the time when flg22 was added. (**D**) Quantitative analysis of peak features for Ca^2+^ traces from N, N+1 and N+2 cells. Each data point in the box plots represents an average value measured from 5–15 cells from a single cotyledon; data from 8 cotyledons from 3 independent experiments denoted by different shapes are presented in each box plot. One-way ANOVA and Tukey’s HSD test, different letters indicate significant differences with P < 0.05.

**Fig. 2.**
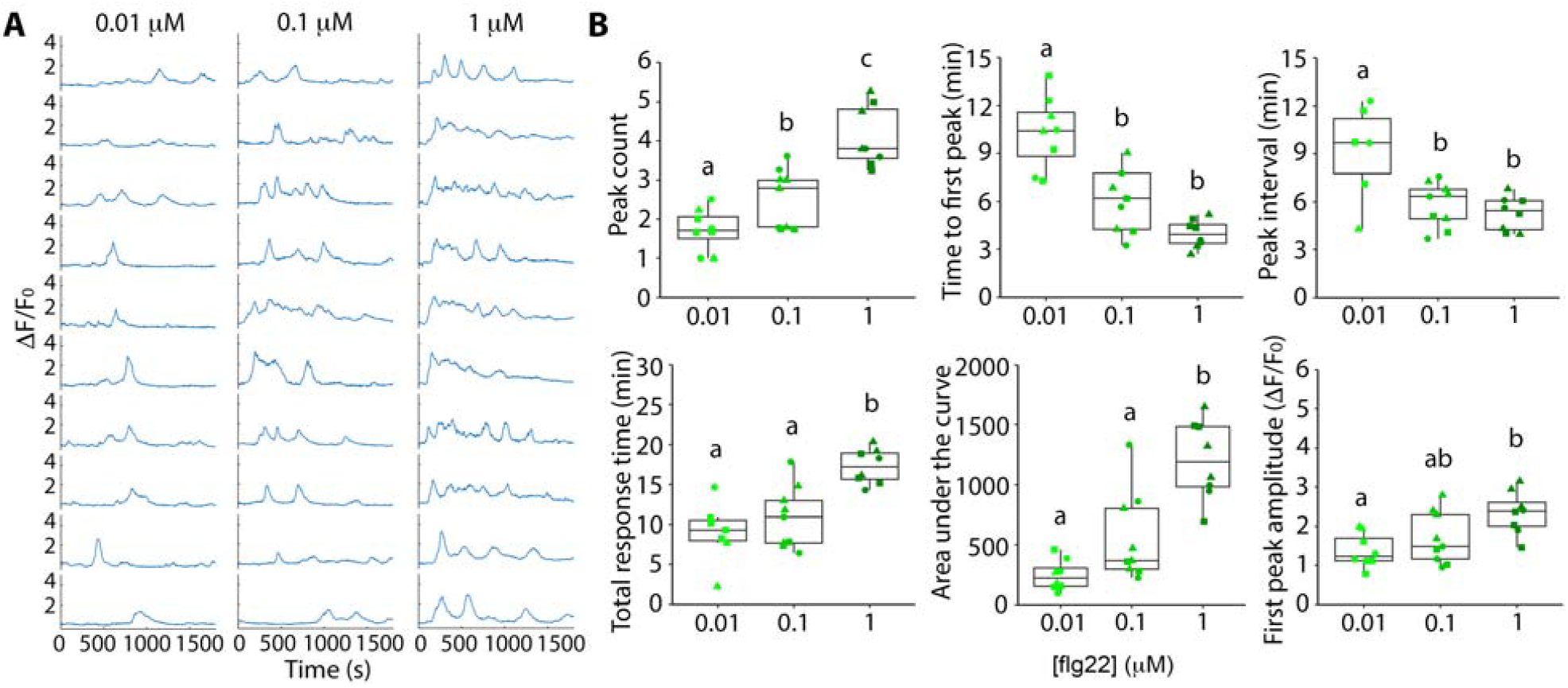
Features of MAMP-induced Ca^2+^ signatures in initiator cells are dose dependent. (**A**) Representative intensity ratio traces from initiator cells following treatment with 0.01, 0.1, or 1 µM flg22. Time 0 represents when flg22 was added to the imaging chamber. (**B**) Quantitative analysis of peak features of Ca^2+^ traces from initiator cells. Each data point in the box plots represents an average value measured from 3–5 initiator cells from a single cotyledon; data from 8–9 cotyledons from 3 independent experiments denoted by different shapes are presented in each box plot. One-way ANOVA and Tukey’s HSD test, different letters indicate significant differences with P < 0.05.

The specific pattern of defense-induced [Ca^2+^]_cyt_ signals at the single-cell level as well as the kinetics of spatial spread are largely unknown. To address this, we analyzed the spatiotemporal dynamics of flg22-induced Ca^2+^ signals across groups of epidermal cells. Interestingly, although flg22 was applied globally and uniformly at a saturating concentration (1 µM; *24*), individual epidermal cells did not show a synchronized response nor a homogenous Ca^2+^ dynamic pattern. Instead, a common spatial pattern was observed: the [Ca^2+^]_cyt_ fluctuations often initiated at a single or a small subset of cells (designated initiator cells), and then a Ca^2+^ wave propagated from the initiator cell(s) to immediately adjacent neighbor cells roughly in a radial pattern (Fig. 1A; fig. S4A; movies S1 and S2). In some cases, a Ca^2+^ wave initiated from a subcellular region within a single cell, whereas in other cases 2 to 3 adjacent cells responded almost simultaneously (fig. S4A; movies S1 and S2). Waves of Ca^2+^ were also observed to initiate from guard cells, however, the frequency of those events was only ∼10% (10 out of 93 waves) of all waves analyzed.

To better understand the heterogeneity of Ca^2+^ signals across epidermal cells, we quantified the Ca^2+^ signature in initiator cells as well as the cells surrounding the initiators. For analysis of Ca^2+^ signals with high throughput and spatial specificity, we utilized LeafNet (*34*), a deep-learning based software tool, to automatically segment the puzzle piece-shaped pavement cells (fig. S1, B and C). Since the cytosolic R-GECO1 signal mainly resides in a thin peripheral layer of the cells when imaged at a medial plane through epidermal cells, the Ca^2+^ intensity profiles were only extracted from the segmented perimeter of each cell (fig. S1C), and the mean fluorescence intensity from individual cells was converted into the ratio ΔF/F_0_ for normalization and statistical comparisons (*27*; fig. S1D). To quantify the Ca^2+^ oscillation pattern and peak features, we developed algorithms using the MATLAB *findpeaks* function to extract peak characteristics, including the number of peaks, peak prominence, peak width at half maximum (WHM), time to 1^st^ peak, mean peak interval, area under the curve (AUC), and total response time (designated as the duration from the 1^st^ peak to the last peak; fig. S1D).

Since our observations suggest that Ca^2+^ signal initiation and propagation exhibit a defined spatiotemporal pattern, with initiation sites often located 2–3 cells apart (fig. S4A), we grouped the epidermal cells into 3 populations based on their spatial proximity to the signal initiation site: initiator cells (N; i.e., epidermal cells with the earliest Ca^2+^ signal increase), cells immediately adjacent to initiator cells (N+1), and cells neighboring the N+1 cells (N+2; Fig. 1B). Statistical analysis of Ca^2+^ signatures among the three groups of cells showed that initiator cells and their close neighbors shared similar spatiotemporal dynamics: both responded to flg22 within 3–4 min, albeit with a slightly delayed time to 1^st^ peak (4.0 ± 0.9 min) in the N+1 cells that was not significantly different from initiator cells (3.7 ± 0.7 min; Fig. 1C, D; fig. S5). Initiator and N+1 cells also showed a similar oscillatory pattern, with an average of ∼4 peaks (at 4–5 min intervals) and a total response time of ∼15 min (Fig. 1, C and D; fig. S5). Although N+1 cells had a slightly reduced number of peaks, total response time, AUC, and increased peak interval, these parameters were not significantly different when compared to those in initiator cells (Fig. 1D). The similarity of Ca^2+^ features between N and N+1 cells suggests a coordinated or dependent relationship in their signaling, likely driven by the propagation and/or diffusion of Ca^2+^ waves between those cells. In contrast, N+2 cells showed a significantly reduced number of peaks (2.5 ± 0.6 peaks), a shortened total response time (11.2 ± 3.8 min), as well as a prolonged time to 1^st^ peak (5.0 ± 1.1 min) when compared to N and N+1 cells (Fig. 1, C and D; fig. S5), suggesting a different pattern or more likely an attenuated response. No significant differences for the AUC or the 1^st^ peak amplitude parameters were detected in N+2 cells when compared to the other two cell groups, however, this was perhaps due to the large variations of peak amplitude among individual cells (Fig. 1D). These results reveal heterogenous Ca^2+^ signaling at the single-cell level as well as spatiotemporally coordinated Ca^2+^ signaling (organized by traveling Ca^2+^ waves) at the multicellular level in response to global pathogen signals.

To test whether the spatiotemporal Ca^2+^ response was specific to a particular MAMP or DAMP, we treated cotyledons globally with chitin, an elicitor from fungal cell walls (*35*), and Pep1, a plant DAMP peptide (*36*), and analyzed the single-cell Ca^2+^ signatures in epidermal cells (fig. S6). Interestingly, the chitin- and Pep1-induced Ca^2+^ signals also exhibited a small subset of cells acting as initiation sites and local traveling wave propagation to a limited number of neighboring cells, resembling the pattern observed for 100 nM flg22-treated tissue (fig. S6, A and B; movies S3 and S4). Initiator cells elicited by chitin or Pep1 also displayed defined oscillatory patterns of [Ca^2+^]_cyt_, with average values of time to the 1^st^ peak, peak interval, total response time, and 1^st^ peak amplitude similar to those observed with flg22 treatment; however, Pep1 treatment resulted in a modest but significant reduction in the number of peaks and AUC (fig. S6, C and D). These results suggest that a small subset of cells act as “first responders”, initiating Ca^2+^ wave spread in the epidermis, is likely a feature common to PTI responses triggered by various types of MAMP or DAMPs. However, the oscillatory patterns of these initiator cells may vary depending on the specific elicitor.

It is unclear how initiator cell(s) are determined in response to global and uniform MAMP treatment. When a large field of view at low magnification was observed, the distribution of initiator cells and sites across the tissue of epidermal pavement cells on a cotyledon appeared to be random (fig. S4A; movie S2), and we did not observe a specific pattern across multiple experiments when similar cotyledon regions were imaged. Whether calcium spiking preceded the oscillations, and thereby defined the initiator cells, was analyzed by characterizing the average number of peaks observed for all three cell types during the first 10 min run-in period. On average, all epidermal types showed approximately less than one peak during the 10 min preceding flg22 addition and no differences were detected between initiator, N+1 and N+2 cells (fig. S4D), suggesting that initiator cells do not have higher Ca^2+^ spiking activity prior to MAMP treatment. To investigate whether the three cell groups correlated with cell size, we measured the cell area from 50–88 cells for each cell group and the results showed that initiator cells were on average larger than the other two cell populations, whereas N+1 and N+2 cells had similar sizes (fig. S4, B and C). Interestingly, this spatial pattern of Ca^2+^ wave spread resembles the *initiator vs. standby* or “leader-follower” pattern described in animal cells for intercellular Ca^2+^ wave propagation (*37, 38*), likely representing a conserved mechanism for coordinated Ca^2+^ signal transduction at the multicellular level.

### The pattern of oscillations in initiator cells depends on flg22 concentration

To investigate whether the spatiotemporal pattern of Ca^2+^ signals is dependent on the dose of elicitor, we treated cotyledons with different concentrations of flg22 and analyzed the single-cell Ca^2+^ signatures. We focused on initiator cells, as the Ca^2+^ dynamics in this cell population were most likely directly related to flg22 perception by the PRR complex. Quantification of peak parameters showed that 1 µM flg22 induced a typical oscillatory pattern of Ca^2+^ signals with an average of 4.1 ± 0.8 peaks (mean ± SD) at 5.3 ± 1.1 min intervals and lasting for 17.1 ± 2.3 min, whereas with lower doses of 0.1 and 0.01 µM flg22 the number of Ca^2+^ peaks was reduced to 2.5 ± 0.8 and 1.7 ± 0.5, respectively (Fig. 2). In addition, the mean peak intervals increased to 5.8 ± 1.4 and 9.1 ± 2.9 min, and the total response time was reduced to 12.2 ± 4.2 and 8.7 ± 3.6 min, respectively, in 0.1 and 0.01 µM flg22-treated cells (Fig. 2B). There was also a dose-dependent reduction of the first peak amplitude and AUC in 0.1 and 0.01 µM treatments when compared to cells treated with 1 µM flg22 (Fig. 2B). These results reveal that the amplitude, frequency, and duration of the oscillatory Ca^2+^ signal at the single cell level are dependent on the dose of elicitor stimulation.

### Intercellular Ca^2+^ waves elicited by MAMPs propagate at a constant speed with limited travel distance

To further understand the mechanism of Ca^2+^ signal transmission at the multi-cellular level, we characterized the intercellular Ca^2+^ wave properties induced by MAMPs. Local spread of Ca^2+^ waves has been demonstrated in Arabidopsis epidermal pavement cells when cells are wounded or mechanically stimulated. Wounding of Arabidopsis leaf epidermal cells with microneedles or laser ablation induces a fast Ca^2+^ wave that propagates in local tissues, and the initial wave speed can reach up to 7–10 µm/s (*10, 11, 13*). To recapitulate the mechanically-induced Ca^2+^ waves and to directly compare it with MAMP-induced Ca^2+^ waves under our experimental set-up, we applied targeted laser ablation to damage a single cell and induce Ca^2+^ wave propagation in the cotyledon epidermis. Consistent with previous reports, immediately following laser irradiation of an epidermal cell, a local [Ca^2+^]_cyt_ elevation was detected and a radial traveling wave propagated from the ablated cell to distal cells (Fig. 3A; movie S5). To estimate the wave speed, we developed an analysis method based on Bellandi et al. (*10*) by extracting the mean fluorescence intensities from a series of concentric rings with the wounded cell set as the center (fig. S7, A and B), since the Ca^2+^ waves traveled in a radial pattern. Analysis of the peaks of the fluorescence traces in concentric rings allowed the wave travel distance to be plotted over time, and therefore the wave speed could be estimated (fig. S7C). Both the kymograph and wave speed analysis showed that the wound-induced traveling wave did not propagate at a fixed rate, instead, the wave exhibited a high initial speed of 4–5 µm/s near the wound site, and the speed gradually diminished as the wave traveled further, until reaching a plateau of ∼1 µm/s before the wave fully dissipated (Fig. 3, B to D). The traveling wave kinetics induced by laser wounding were consistent with a previous report of needle wound-induced local Ca^2+^ waves, and the same study demonstrated that the wound-induced Ca^2+^ wave progression coincided with apoplastic diffusion of glutamate that was released from the wounded cell, acting as a DAMP signal to induce Ca^2+^ influx and wave propagation by activating glutamate receptor-like (GLR) channels (*10*). Since wound-induced Ca^2+^ signals are mediated by apoplastic diffusion of amino acids, we further confirmed this by treating cotyledons globally with a dose series of glutamate. Glutamate treatment at 10 mM or above induced an immediate and synchronized Ca^2+^ burst among most epidermal cells, without the typical pattern of initiator cells or sites leading the wave in MAMP-treated cells (fig. S8). In addition, glutamate triggered only a single Ca^2+^ spike in most of the cells analyzed, rather than the oscillations observed after MAMP treatment (fig. S8, B and C), indicating a distinct Ca^2+^ signature compared to those induced by PRR perception of MAMPs.

**Fig. 3.**
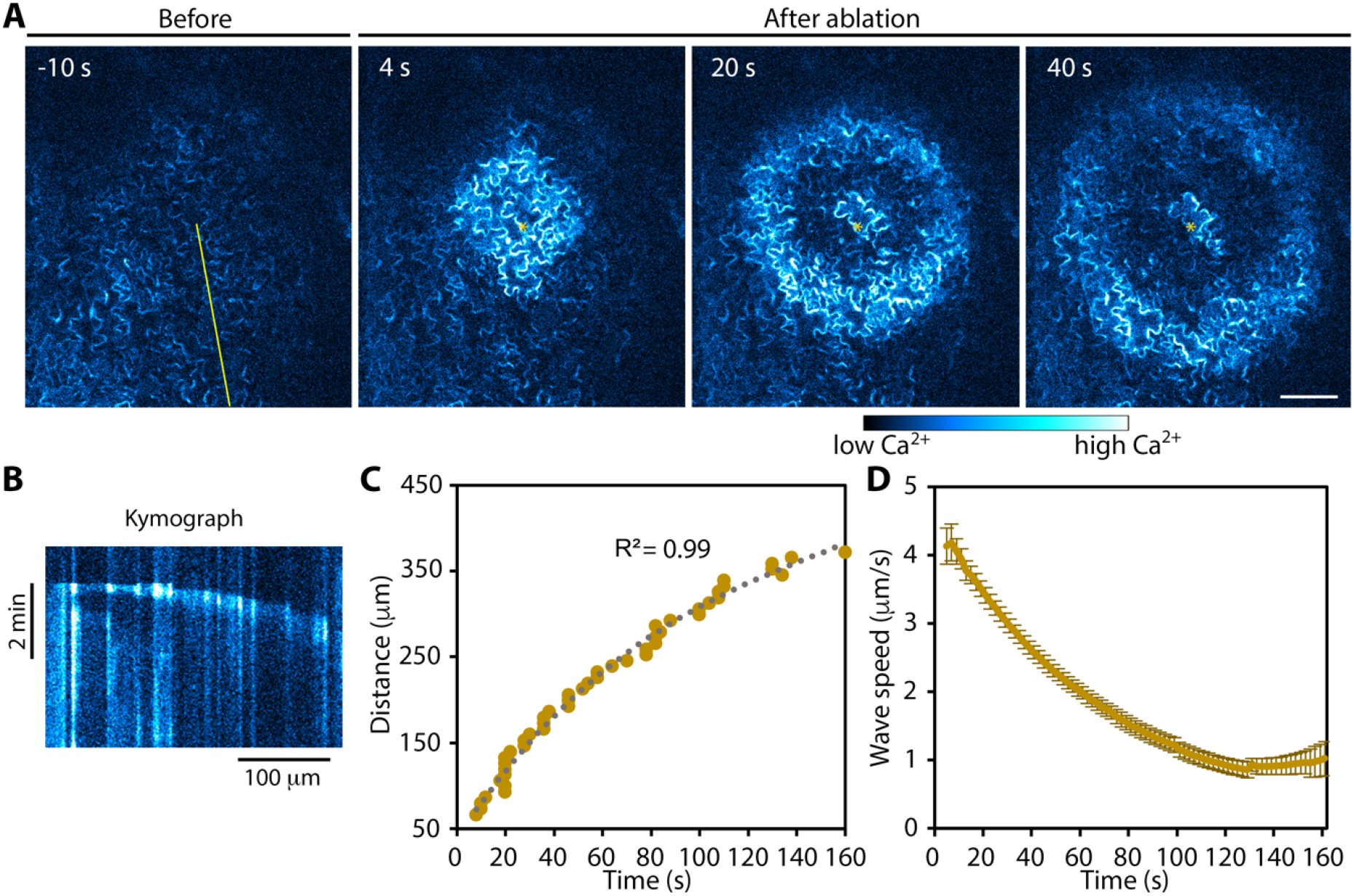
Wounding of a single epidermal cell with laser irradiation induces a traveling wave of Ca^2+^ that spreads by diffusion. (**A**) Representative timelapse image series shows the laser ablation-induced Ca^2+^ wave propagation in cotyledon epidermal cells expressing R-GECO1. See also Supplemental Movie S5. The ablated cell is marked with an asterisk. Bar = 100 µm. (**B**) Representative kymograph generated from the yellow line drawn in (A). The trajectory of the wave front shows a non-linear curve suggesting that speed varies as the wave progresses. (**C**) A representative plot of the wave travel distance over time using the method shown in Figure S7. The curve was fitted with a polynomial of degree 3 (dashed line). (**D**) The first derivative of the polynomial (as shown in C) was calculated as the speed of the wave front over time. Values given are means ± SE (n = 11 cotyledons from 3 independent experiments).

To further compare MAMP-induced Ca^2+^ wave spread pattern with those induced by wounding or glutamate, we analyzed the MAMP-induced Ca^2+^ wave speed with the ring scan method described in fig. S7, since the waves propagated from the initiation sites in a roughly radial pattern (Fig. 4, A and B; movie S6), and this spread could be fit with a series of concentric rings (fig. S7A). Interestingly, in the majority of waves analyzed, the plot of distance traveled over time or kymographs could be fitted with a linear curve (Fig. 4C; fig. S7, C and F), suggesting that the wave traveled at a constant speed. Among 93 waves analyzed from 32 cotyledons, 75 could be fitted with a linear curve (80.6%). In the rest of the cases, 11 waves (11.8%) showed an initial delay in propagation, suggesting a slower speed at the early phase, whereas 7 waves (7.5%) showed a faster speed and then slowed down as they traveled further (fig. S7, D and E). Analysis of fluorescence intensity profiles at various positions along the traveling wave (origin, half, and end of the wave’s path) revealed a reduction of signal amplitude and an increase of signal half-maximal width as the wave propagated to the distal end (Fig. 4, E to H). Estimation of traveling wave speed from the linear fit showed that the waves traveled at 1–1.5 µm/s when 1 µM flg22 was applied to epidermal tissue (Fig. 4D), which is 4–5 times slower than the initial wave speed induced by laser ablation or wounding. The traveling wave speed was also dependent on the concentration of flg22, as the average speed was reduced to 0.9 ± 0.2 µm/s (mean ± SD) in 0.01 µM flg22-treated cotyledons compared to 1.3 ± 0.2 µm/s with 1 µM flg22 treatment (Fig. 4D). Analysis of chitin- and Pep1-induced Ca^2+^ waves showed that they also traveled at a constant velocity, although chitin-induced waves were significantly slower (0.9 ± 0.1 µm/s), when compared to 1.2 ± 0.3 µm/s and 1.3 ± 0.2 µm/s for flg22- and Pep1-treated cells, respectively (fig. S6E). The slow yet constant travel speed suggests that MAMP-induced Ca^2+^ spread is likely driven by an active propagation mechanism, rather than by symplastic or apoplastic diffusion.

**Fig. 4.**
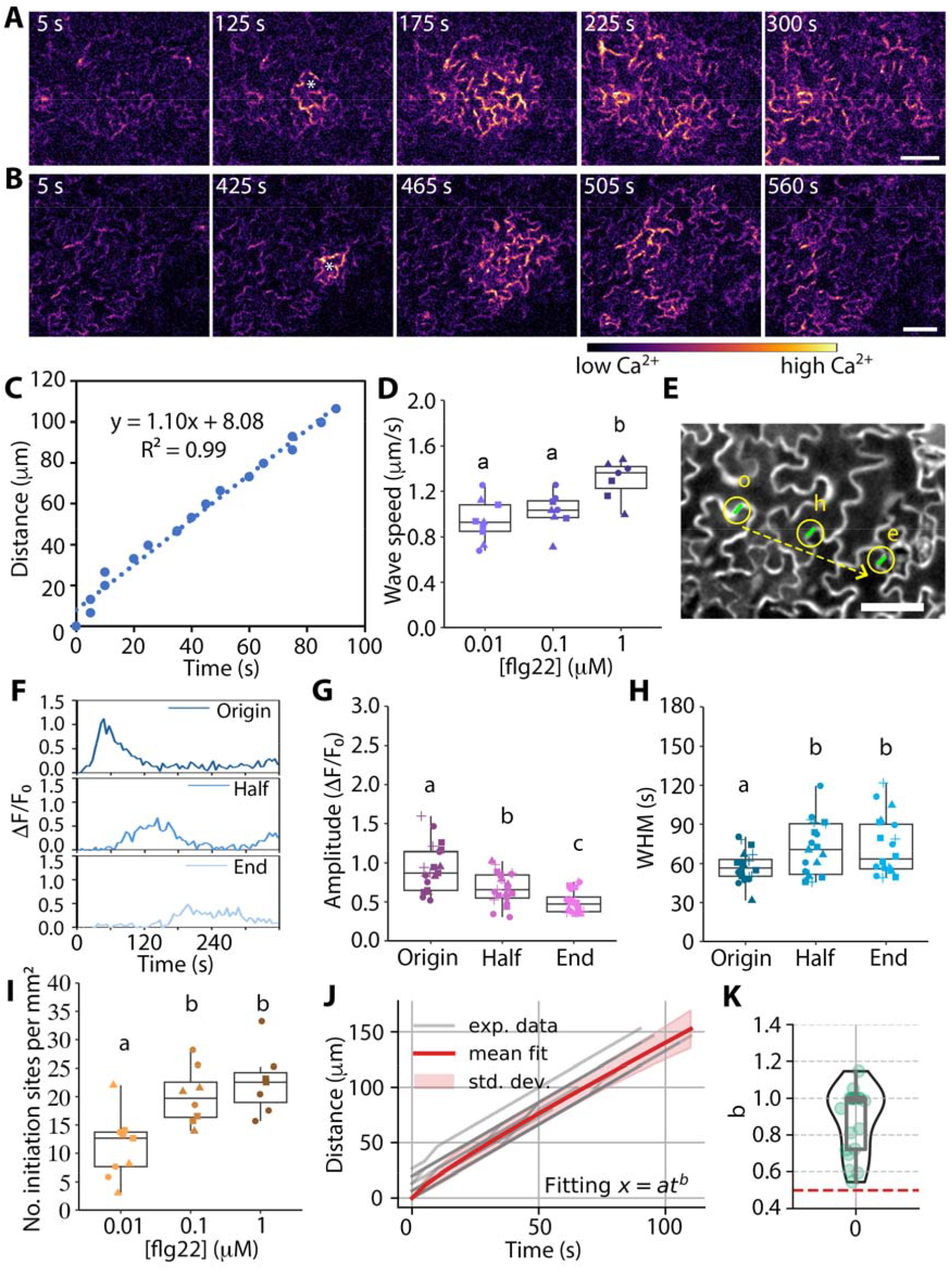
Traveling waves of Ca^2+^ elicited by flg22 propagate at a constant speed and have a limited area of spread. (**A, B**) Two representative timelapse image series show a traveling wave of Ca^2+^ propagating across cotyledon epidermal cells. See also Supplemental Movie S6. Cotyledons in (A) and (B) were treated with 0.1 µM or 0.01 µM flg22, respectively. Initiator cells are marked with an asterisk. Bars = 50 µm. (**C**) A representative plot of wave travel distance over time with a linear function fitted to estimate wave speed (see Figure S7 for methods). Time = 0 represents the initiation of a wave and the same applies to subsequent panels. (**D**) Quantitative analysis of Ca^2+^ wave speed from cotyledons treated with 0.01, 0.1, or 1 µM flg22. (**E**) A 10-µm long line (green) near a wave origin (o), half (h), and end (e) of the travel distance was selected as region of interest for extraction of Ca^2+^ intensity profiles and analysis of peak features. The dashed arrow marks the direction of wavefront travel. Bar = 50 µm. (**F**) Representative Ca^2+^ intensity profiles at the origin, half and end of the travel distance of a wave as shown in (E). (**G, H**) Quantification of calcium signal amplitude (G) and width at half maximum (WHM; H) at three wave locations as shown in (E) and (F) from cotyledons treated with 100 nM flg22. Data points in the box plots represents individual waves; data from 21 waves from 4 independent experiments denoted by different shapes are presented in each box plot. (**I**) Quantification of the density of initiation sites for traveling waves in the cotyledon epidermis tissue following treatment with 0.01, 0.1, or 1 µM flg22. Each data point in the box plots represents an average value measured from 3–6 waves from a single cotyledon; data from 8–9 cotyledons from 3 independent experiments denoted by different shapes are presented in each box plot. One-way ANOVA and Tukey’s HSD test, different letters indicate significant differences with P < 0.05. (**J**) Plot showing distance (x) traveled by the Ca^2+^ wavefront with respect to time (t) for 20 different waves from cotyledons treated with 100 nM flg22. Each dataset was independently fit with a power-law model of the form *x=at*^*b*^. The plot shows averaged fit across samples, with shaded regions representing the standard deviation of the fits. (**K**) A combination of box and violin plot visualizes the distribution of power-law exponent (*b*) across samples shown in (J). The dashed red line indicates *b* = 0.5, which corresponds to the expected value for a diffusion-driven process.

In addition, the distance traveled by MAMP-induced waves appeared to be limited; usually limited to 100–200 µm, or about 3–4 cells away from the wave initiation site (Fig. 4, A and B). This was in contrast to wound-induced waves, which typically traveled further, with a linear distance of 350–400 µm, or about 6–8 cells away from the wound site (Fig. 3, A and C). The individual wave travel distance was only estimated in cotyledons treated with low doses of MAMPs where there was no merging of waves with each other and the wave edges could be clearly determined, as high-dose treatment often resulted in a high density of initiation sites within the epidermal layer, consequently leading to the merging of wave fronts that propagated over substantially longer distances.

To determine the density of wave initiation sites in cotyledon epidermal cells, we quantified the total number of initiation sites in the imaged field (681 × 681 µm^2^). The results showed that there was an average of 11.2 ± 5.6 initiation sites per mm^2^ in cotyledons treated with 0.01 µM flg22, whereas this density significantly increased to 20.1 ± 4.9 and 22.5 ± 5.8 sites/mm^2^ for 0.1 and 1 µM treatments, respectively (Fig. 4I). No significant difference of initiation site density was detected between 0.1 and 1 µM treatments, suggesting that the total number of initiation sites or waves that can be elicited in the cotyledon epidermis is limited. In addition, a similar density of initiation sites was observed in cotyledons treated with chitin and Pep1 compared to that with flg22 (fig. S6F), indicating that the number of initiation sites is not dependent upon the type of elicitor.

Overall, our results demonstrated that PRR perception of various MAMPs/DAMPs induces a similar type of Ca^2+^ wave that propagates at a constant speed but over a limited distance, which is distinct from the waves induced by touch or wounding.

### Calcium-induced calcium release (CICR) is sufficient to drive slow traveling waves with a constant speed

To identify possible mechanisms for MAMP-induced Ca^2+^ wave propagation, we first tested whether Ca^2+^ release at an initiation site could drive wave transmission purely through diffusion. To test this hypothesis, a power law model of the form *x = atb* was fit to the experimental data for 100 nM flg22 treatments, where *x* represents the distance traveled by the wavefront and *t* represents time. For processes regulated by diffusion, the distance traveled by a wave is expected to scale with the square root of time (*b* = 0.5). However, our analysis of multiple independent wavefronts elicited by treatment with 100 nM flg22 revealed that the average power-law exponent, *b*, was greater than 0.5 (Fig. 4, J and K). This suggests that wave propagation is not governed by simple diffusion. Additional mechanisms involving Ca^2+^ signal amplification must be involved for driving experimentally observed Ca^2+^ wave patterns.

One conserved mechanism for Ca^2+^ wave transmission is calcium-induced calcium release (CICR), a process described in a wide variety of eukaryotic cells for active propagation of Ca^2+^ signals (*39, 40*). Our observation that although the tissue was treated globally with MAMP elicitors, traveling waves of Ca^2+^ consistently followed a sequential time order from initiator cells to neighboring cells and transmitted at a constant speed, supports a model that the wave is propagated by Ca^2+^ itself. To test this hypothesis and describe the traveling wave mathematically, we used the fire-diffuse-fire model developed by Dawson et al. (*41*) to simulate CICR-driven wave propagation based on the empirical data from our flg22-treated dose response studies (fig. S9). For more details on the modeling, refer to the Methods section. Although CICR generally refers to a process in which calcium induces further calcium release from intracellular stores, in our model we did not specify the types of channels or stores, as it remains unclear which Ca^2+^ channels might play such a role in PTI or whether they are located on the PM or membranes of other intracellular stores (*21*).

Our primary aim was to test whether CICR alone could reproduce the experimentally observed constant wave speed. To achieve this aim, we first performed Latin Hypercube sampling to generate 20,000 parameter sets (*42*), varying model parameters within biologically relevant ranges (*14, 41*). We set the Ca^2+^ diffusion coefficient, *D*, to 20 μm^2^/s, a value commonly used in the literature (*14*). Our model also uses the assumption that cell-to-cell diffusion of Ca^2+^ is not restricted by plasmodesmata (*14*). For each estimated parameter set (Fig. 5, A and B), we calculated the average wave speed and the R^2^ value of a linear fit for wavefront distance versus time (Fig. 5C). Parameter sets that produced constant wave velocities within the experimental range defined by the 1 μM flg22 treatment dataset (mean = 1.3 ± 0.2 µm/s; range = 0.89 – 1.76 µm/s) and high R^2^ values (> 0.99) were selected for further analysis (Fig. 5, A and B). 22 out of 20,000 parameter sets passed these selection criteria. Our results demonstrate that a simple CICR mechanism was sufficient to replicate the constant wave speed of ∼1.3 µm/s, with parameters that are within a reasonable biological range (Fig. 5; table S1).

**Fig. 5.**
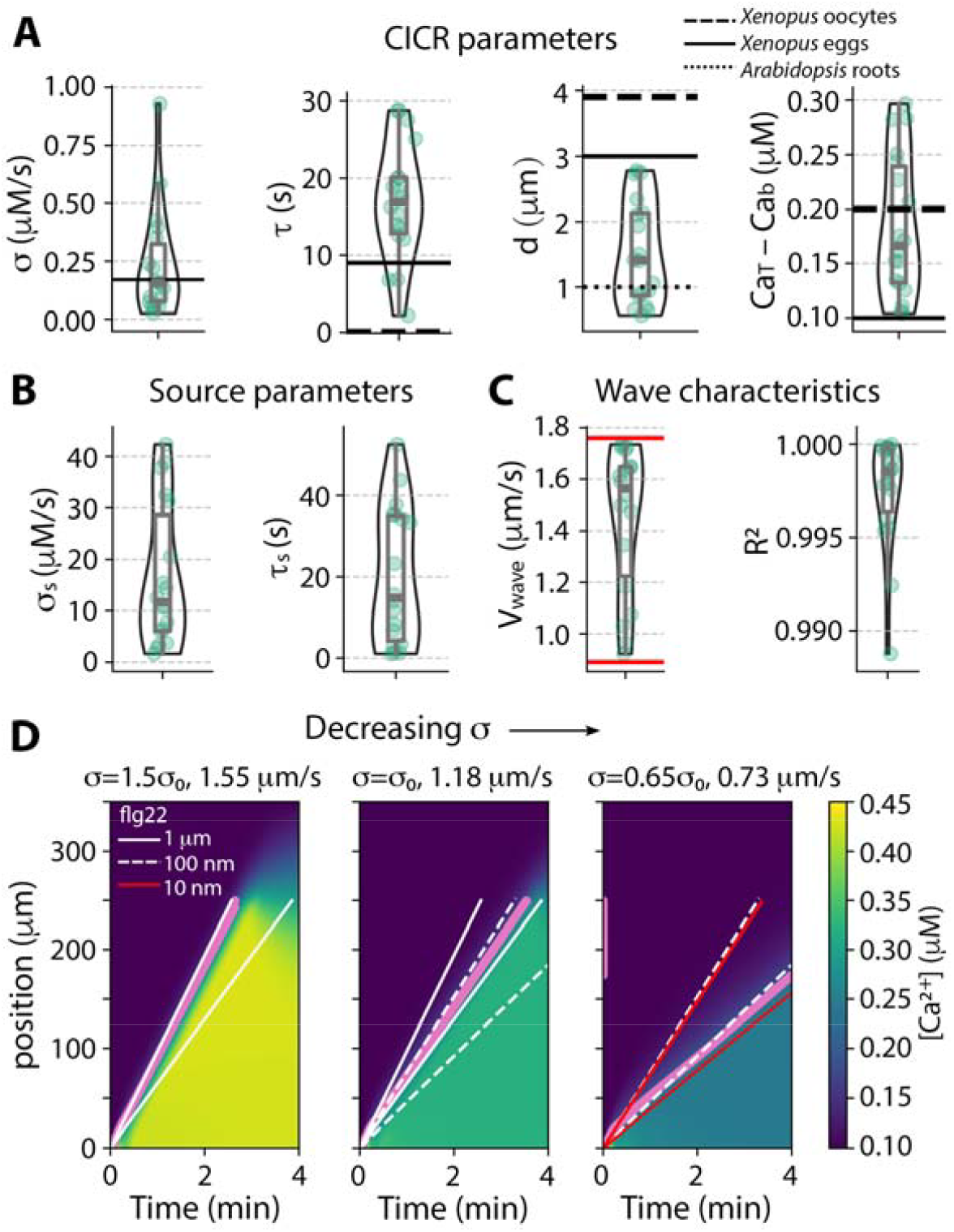
A mathematical model for calcium-induced calcium release is sufficient to produce constant wave speed consistent with empirical data. Empirically-determined wave speeds from 1 µM flg22 treatment, as shown in Figure 4D, were used to calibrate the 1D calcium-induced calcium release (CICR) model. (**A, B**) A combination of box and violin plots illustrates the distribution of parameter values in the calibrated model. The box represents the interquartile range (IQR), whereas the whiskers extend to 1.5 times the IQR. The horizontal line within the box marks the median. The black solid and dashed lines indicate the reported parameter values from other studies (*14, 41*). (**C**) A similar visualization illustrating wave speed (*ν* _wave_) and R^2^ values for fitted parameter sets. Red lines indicate the maximum and minimum wave speeds measured from 1 μM flg22 treatments that were used for calibration. (**D**) Kymographs showing changes in Ca^2+^ wave dynamics when the parameter σ, representing the rate of flux of Ca^2+^ through the channel, was varied. The y-axis represents spatial location, whereas the x-axis represents time. Here, parameter σ was increased by 50% (left) and decreased by 35% (right) for a representative parameter set. Pink points mark the activation times for each channel location, representing the model-predicted leading edge of the traveling wave. Corresponding wave edges of the experimental data for different concentrations of flg22 are shown as mean ± SD.

Among these parameters, we observed that the parameter *σ*, representing the rate of Ca^2+^? entry upon CICR channel activation, was inversely proportional to the channel activation time constant, *τ* (fig. S10C). Moreover, channels with lower *σ* and higher *τ* values produced waves with lower speeds. Additionally, we found that higher distance between Ca^2+^ channels, *d*, required a higher *σ* to maintain constant wave speed within the desired range.

Within our model, we assumed that both flg22 perception and Ca^2+^ are necessary to activate CICR channels. Consequently, a reduction in flg22 concentration could be represented in the mathematical model by decreasing *σ*. Reducing the flg22 concentration from 1 μM to 10 nM resulted in a 30.4% decrease in average wave speed for the empirical data (Fig. 4D). Using a single selected parameter set, we varied *σ* to validate these findings, observing that a 35% reduction in *σ* led to a 39.8% decrease in wave speed (Fig. 5D). Across different estimated parameter sets, decreasing σ by up to 30% consistently reduced average wave speed (fig. S10A). Conversely, increasing *σ* raised the wave speed (Fig. 5D). In summary, our results demonstrate that a simple CICR mechanism is sufficient to replicate the constant wave speeds determined empirically and recapitulates the flg22 concentration dependence of wave speed.

We observed that MAMP-induced Ca^2+^ waves traveled only short distances and diminished in amplitude over time (Fig. 4; fig. S7), suggesting that a robust signal attenuation mechanism operates simultaneous with CICR and limits the wave transmission distance as well as perhaps the wave speed. To test this, we introduced a sink to our mathematical model and assumed it was activated by Ca^2+^ binding. This assumption represents known biological sinks like the autoinhibited Ca^2+^-ATPase (ACA) pumps in plants (*31, 43, 44*) or SERCA pumps in animal cells (*45*). We modeled the rapid binding of Ca^2+^ to these pumps with Hill functions (see Methods for details). Using a single parameter set from the previous best fits, we varied the parameter *V* _*sink,max*_, representing the maximum flux of Ca^2+^ through the sink. Increasing *V* _*sink,max*_ slowed wave propagation and led to signal attenuation (Fig. 6A; fig. S11B (left column)). This increase in *V*_*sink,max*_ also led to a slight deviation from a constant velocity profile during wave propagation, as reflected in decreased R^2^ values. We performed similar analyses across all parameter sets that produced constant wave speeds within biologically relevant limits (Fig. 5A), with a sink added to each parameter set (fig. S11, E to G).

**Fig. 6.**
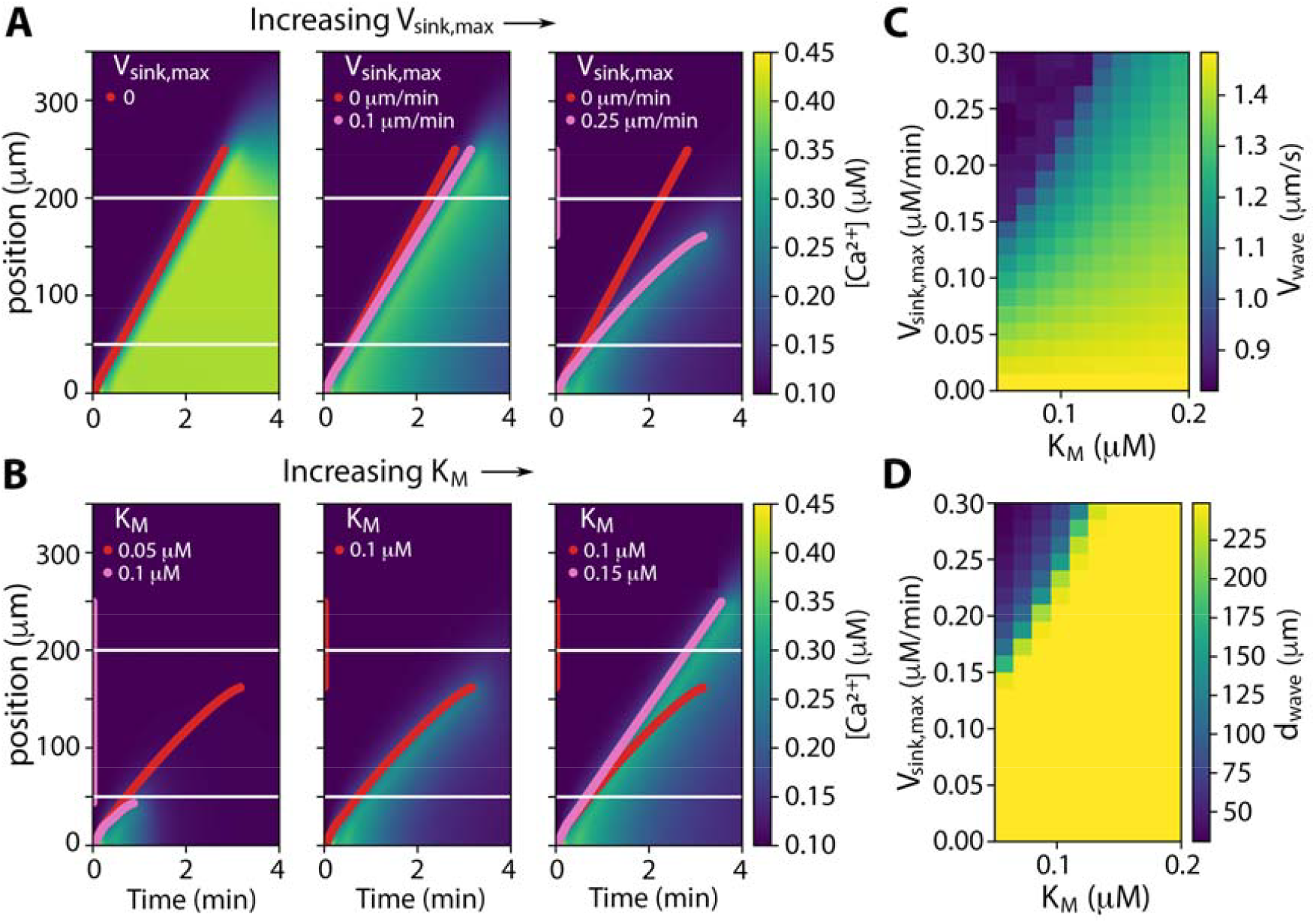
Adding a term for a sink, representing a Ca^2+^-dependent pump, to the mathematical models allows for attenuation of the traveling wave. (**A**) Kymographs depicting spatiotemporal changes in Ca^2+^ concentration in response to increasing the parameter *v*_*sink,max*,_ with Km held constant at 0.1 µM. Pink points mark the activation times for each channel location, representing the model-predicted leading edge of the traveling wave when a sink was added to the model. Red line shows the model-predicted leading edge of the traveling wave without a sink. (**B**) For *V* _*sink,max*_ = 0.25 μM/s, the parameter *K*_*M*_ was adjusted by increasing (right) and decreasing (left) it by 50%. The leading edge of the traveling wave is color coded as red for the base case (middle, *K*_*M*_ = 0.1 µM) and overlaid on top of the leading edges in pink when *K*_*M*_ was varied (left and right). Horizontal lines in the kymographs mark the experimentally observed bounds for the distance traveled by Ca^2+^ waves for 1 µM flg22 treatment. (**C, D**) Parameters *K*_*M*_ and *V*_*sink,max*_ were varied within specified ranges. For each parameter combination, we calculated the wave speed (C) and the distance traveled by the wave (D), which are represented as heatmaps.

Adding a sink with *V*_*sink,max*_ = 0.2 μM/min and *K*_*M*_ = 0.1 μM caused reductions in both wave speed and the distance travelled by the wave across all the biologically relevant parameter sets.

For the selected *V*_*sink,max*_ value that caused signal attenuation, we further varied *K*_*M*_, which reflects the Ca^2+^ concentration for half-maximal activation of pumps (Fig. 6B; fig. S11B (right column)). Decreasing *K*_*M*_ reduced both the distance travelled and wave speed, whereas increasing *K*_*M*_ had the opposite effects. We also quantified the combined effects of *V*_*sink,max*_ and *K*_*M*_, finding that higher *V*_*sink,max*_ and lower *K*_*M*_ values led to slower waves that travelled short distances before attenuating (Fig. 6, C and D; fig. S11A).

To further compare the modeling results with the experimentally observed Ca^2+^ wave dynamics, we analyzed Ca^2+^ signals at three CICR channel locations along the 1D wave path, representing origin, half and end of the travel distance (fig. S11D).

Analysis results revealed that across all parameter fits with an added sink (*V*_*sink,max*_ =0.2 μM/min and *K*_*M*_ = 0.1 μM), the half-maximal width of the wave was broader at the wave’s central region compared to the origin and distal ends (fig. S11, H and I). These results were consistent with the analysis of the experimental data for 100 nM flg22 treatment (Fig. 4, E to H). In summary, our modeling results demonstrate that adding a sink facilitates the return of cytosolic Ca^2+^ to basal levels, improving alignment with experimentally observed Ca^2+^ dynamics. The sink also effectively regulates wave speed and limits wave travel distance before it attenuates.

To test our mathematical model predictions that CICR is sufficient to drive slow Ca^2+^ waves with a constant velocity, we experimentally investigated the role of cyclic nucleotide-gated ion channels (CNGCs). The PM-localized CNGCs, CNGC2 and CNGC4, are required for Ca^2+^ influx induced by several MAMPs or DAMPs, including flg22, elf18, chitin and Pep1 (*46*). CNGCs are also involved in propagation of systemic Ca^2+^ waves induced by wounding or high light treatment (*15, 47*). To test whether CNGC2/4 might be the channels responsible for Ca^2+^ influx as well as CICR-driven wave propagation in our system, we quantified the single-cell Ca^2+^ signatures as well as the intercellular wave travel speed induced by flg22 in a *cngc2/4* double mutant (*46*) expressing R-GECO1. Surprisingly, when compared to a wild-type sibling line, neither the Ca^2+^ oscillations in initiator cells or the traveling wave kinetics were affected in *cngc2/4* (Fig. 7, A and B). The *cngc2/4* mutant also had a similar number of wave initiation sites compared to that in wild-type seedlings (Fig. 7C). Our results indicate that CNGC2/4 are not critical for flg22-induced Ca^2+^ influx in cotyledon epidermal cells. It is possible that other CNGC isoforms might play a redundant role in these cells or an unknown type of channel could be responsible for the slow traveling wave transmission observed in this tissue.

**Fig. 7.**
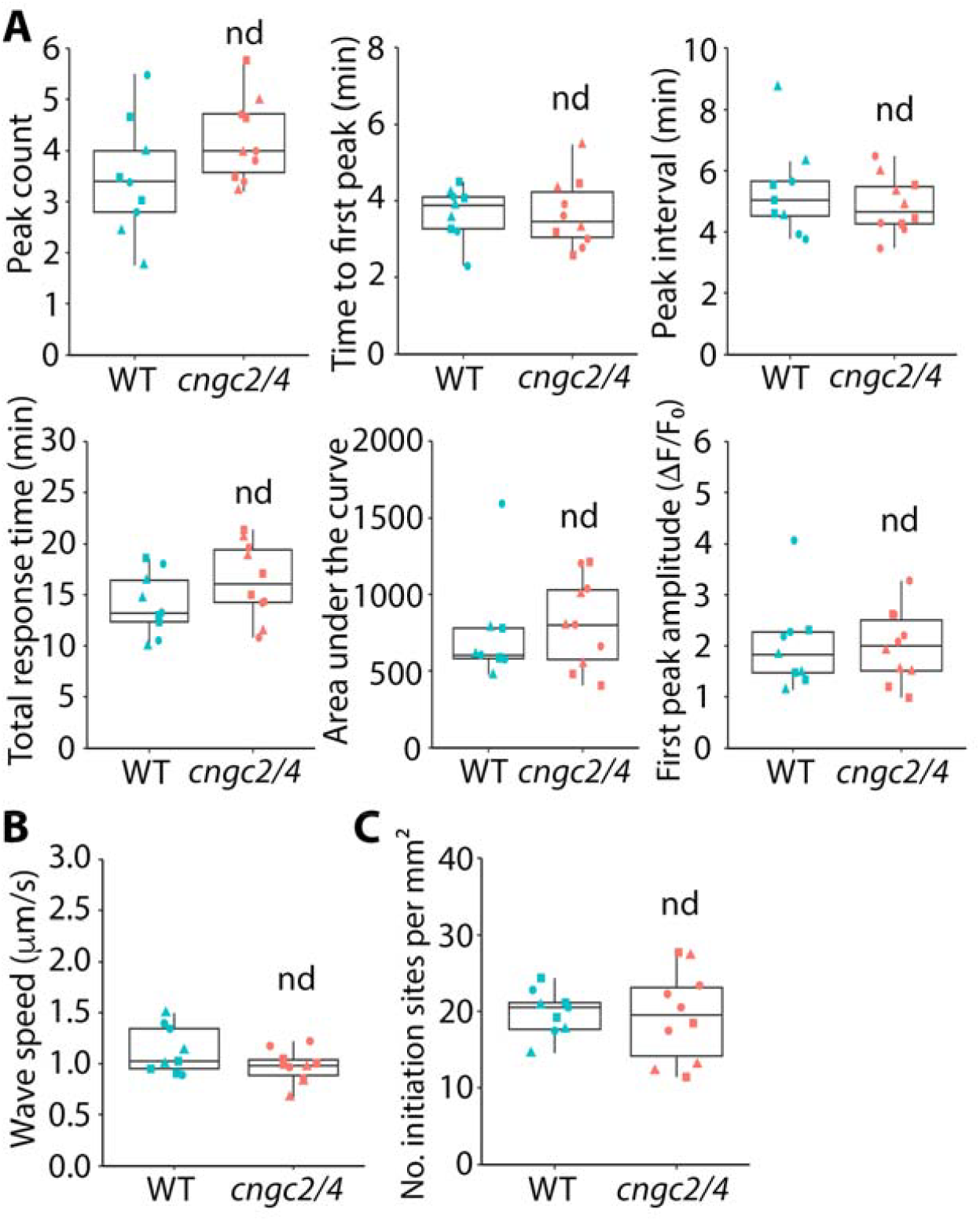
Ca^2+^ signatures and traveling wave speed elicited by flg22 are unaltered in *cngc2/4*. (**A**) The *cngc2/4* double mutant and a wild-type (WT) sibling line expressing R-GECO1 were treated with 100 nM flg22 and the peak features from initiator cells were quantified. (**B**) Quantitative analysis of traveling wave speed in the cotyledon epidermis from WT and *cngc2/4*. (**C**) Quantification of the density of wave initiation sites in the cotyledon epidermis tissue from WT and *cngc2/4*. Each data point in the box plots represents an average value measured from 3–5 cells or 3–6 waves from a single cotyledon; data from 9–10 cotyledons from 3 independent experiments denoted by different shapes are presented in each box plot. Student’s T test, nd: P > 0.05.

## DISCUSSION

Cytosolic Ca^2+^ signatures with distinct spatiotemporal patterns are believed to encode critical information and enable multicellular organisms to respond to specific environmental stimuli with precise outcomes in terms of magnitude, duration and spatial range of the response. Traveling waves of cytosolic Ca^2+^ serve as an essential means of communication and signal propagation between cells. However, the cellular and molecular mechanisms underlying the generation of these stimulus-specific Ca^2+^ signals and traveling waves remain largely unknown, particularly in plant responses to biotic or abiotic stresses. In this study, we characterized Ca^2+^ oscillations and local traveling waves at single-cell resolution during the first layer of plant immune response, pattern-triggered immunity, and examined their specificity upon recognition of various pathogen-related chemical or mechanical signals. Our findings shed light on the molecular mechanisms of plant defense-related Ca^2+^ signaling at both the single cell and multicellular level, with implications for understanding how plants spatially fine-tune immune responses.

Specifically, our findings revealed a distinct mechanism for the generation and transmission of Ca^2+^ traveling wave transmission across a tissue, the epidermis of cotyledons, in response to chemical *vs*. mechanical stimuli that are related to plant defense. We demonstrated that PRR recognition of several MAMPs or DAMPs induced a similar spatial propagation pattern of Ca^2+^ waves that travelled with a constant speed of ∼1 µm/s across a limited number of epidermal pavement cells. In contrast, waves induced by single-cell mechanical damage exhibited a diffusion-like spread and decay pattern with an initial speed of 4–5 µm/s. These results likely reflect two distinct mechanisms for propagation of intercellular Ca^2+^ signals: one driven by apoplastic diffusion of channel-activating molecules, such as glutamate, and the other by a symplastic reaction-diffusion mechanism such as calcium-induced calcium release (CICR). Mathematical modeling supports that a fire-diffuse-fire model, describing a CICR mechanism with a sink, is sufficient to recapitulate the features of the slow and local traveling wave induced by global MAMP treatment.

Traveling waves of Ca^2+^ are best characterized in plant systemic signaling, where they propagate at speeds of several hundred µm/s through the vascular tissue (*7, 48*). How Ca^2+^ signals propagate intra-or intercellularly in local tissues, such as the leaf or cotyledon epidermal pavement cells—which often serve as the first line of external signal perception and transduction—remains largely unexplored. Most studies on local Ca^2+^ traveling waves have focused on mechanical stress responses; for example, touch or wounding of a single leaf epidermal cell or trichome in Arabidopsis induces Ca^2+^ waves that propagate in a radial or ring-like pattern from the site of stimulation (*10, 12, 13*). The reported wave velocities range from 1–10 µm/s and exhibit a diffusion-like spread and decay pattern when measured at high temporal resolution (*10, 13*), consistent with our findings following laser ablation of single cotyledon epidermal cells in this study. A slow (1.63 µm/s) traveling wave of Ca^2+^ was also reported in Arabidopsis roots when a single epidermal cell was damaged by laser irradiation (*49*); however, the temporal resolution did not allow for a full description of the kinetics. By simultaneously imaging Ca^2+^ and glutamate with an apoplastic glutamate-sensing fluorescent reporter, Bellandi et al. (*10*) demonstrate that touch-or wound-induced local Ca^2+^ waves co-propagate with an apoplastic glutamate wave, and mathematical modeling and fitting of the wave curve with a diffusion equation support a simple diffusion-driven model for wave propagation.

Ligand binding to plasma membrane receptors, or the perception of chemical signals like MAMPs, DAMPs and cell wall integrity-related elicitors (*50*), represents a key signaling mechanism in all types of plant cells. However, the molecular mechanisms that underpin Ca^2+^ signal propagation or transmission to neighboring cells after MAMP or DAMP perception are poorly characterized in comparison to those induced by mechanical stimulation. This is likely due to the technical challenges associated with applying local chemical treatments to a single cell or a defined region of a tissue. Our study is the first to characterize a local traveling wave of Ca^2+^ induced by MAMP perception resolved at high spatiotemporal resolution. Global treatment of excised Arabidopsis cotyledons with a variety of MAMPs or DAMPs leads to a qualitatively similar outcome in epidermal pavement cells. Hotspots of signal perception, so-called initiation sites or cells, undergo subcellular and cellular Ca^2+^ oscillations that were quantitatively described. The first peak (and often the second) spreads to neighboring cells as a traveling wave that moves at constant speed (1.3 ± 0.2 µm/s for 1 µM flg22 treatment) but only propagates for a limited distance of 100–200 µm (in low-dose treatments). Quantitative analysis and mathematical modeling of the wave kinetics revealed a distinct propagation mechanism compared to waves induced by mechanical stimulation. The constant wave speed could be recapitulated with a fire-diffuse-fire or CICR model, based on parameters that are within reasonable biological ranges for plant cells. Moreover, a sink or Ca^2+^ pump was required in the mathematical model to convert a Ca^2+^ flood into a wavefront with a trailing edge as well as to limit the distance of transmission.

In animals and several other eukaryotic systems, CICR represents a key mechanism for generating Ca^2+^ transients and waves, and is primarily mediated by Ca^2+^-gated ion channels located on intracellular organelles like the endoplasmic reticulum (*39, 40*). In contrast, CICR mechanisms in plants are less understood, largely due to the absence of direct homologs like the inositol trisphosphate receptors (IP3Rs) or ryanodine receptors (RyRs), which are key players of CICR in animals (*2*). Plants could operate a similar CICR mechanism through the vacuole membrane-localized channel protein Two-Pore Channel 1 (TPC1), which is activated by cytosolic calcium increases (*51*). TPC1 plays a key role in propagating long distance traveling waves of Ca^2+^ induced by salt stress in roots (*14, 52*). However, traveling waves mediated by TPC1 propagate at 400 µm/s in root cortex and endodermal cells, which is hundreds of times faster than the local slow waves we and others observe (*52*). In addition, TPC1 does not play a role in the total averaged Ca^2+^ signal induced by MAMPs when measured in leaf discs (*53*), making it an unlikely candidate for the channel responsible for CICR during MAMP signaling.

Cyclic nucleotide-gated ion channels (CNGCs) are implicated in MAMP-induced Ca^2+^ release and plant immunity (*46*). However, our results indicated that at least for the cell type studied here, 7-day-old epidermal cells from cotyledons, CNGC2 and CNGC4 were not major players for Ca^2+^ release and wave propagation upon MAMP perception. Similarly, other studies also question the role of CNGC2/4 in flg22-induced Ca^2+^ elevations or ROS bursts when whole leaves or seedlings are measured (*54, 55*). Nonetheless, we cannot rule out the possibility that other CNGC isoforms might play a redundant role in these cells or that there are tissue-or development-specific expression patterns utilized by CNGC2/4. Given that CNGCs have been shown to play an indispensable role in propagation of systemic Ca^2+^ waves induced by wounding or high light treatment (*15, 47*), and that their activity can be positively regulated by Ca^2+^-CaM (*56*), they are still strong candidates for the CICR channels that support Ca^2+^ wave propagation in plant cells.

Recent breakthroughs indicate that PTI-induced Ca^2+^ signals could involve multiple classes of ion channels besides CNGCs, including clade 2 GLRs, reduced hyperosmolality-induced Ca^2+^ increase (OSCA) channels, and annexins (*46, 57–59*). In addition, several studies indicate that inositol polyphosphate (InsP)-induced Ca^2+^ release from intracellular stores (IICR) also plays a role in MAMP signaling or plant defense (*60, 61*), especially for flg22 perception (*28, 62*). However, the specific channels activated by InsPs and their subcellular locations remain unknown. Our study raises the possibility that one of these channel classes, or another yet to be identified channel, could function in a CICR-like mechanism to amplify and propagate the Ca^2+^ signals both intra- and intercellularly. Although none of these PTI-related channels are known to be directly activated or positively regulated by cytosolic Ca^2+^ elevations, it is possible that some possess unconventional Ca^2+^-binding sites on their cytosolic surface or are regulated indirectly through Ca^2+^-binding proteins.

Although the constant wave speed could be recapitulated mathematically with a CICR model, we cannot rule out other mechanisms that might generate or assist the propagation of traveling waves in epidermal pavement cells. Although the wave is unlikely to be driven solely by diffusion, it remains possible that molecules released or produced from initiator cells, such as amino acids, other DAMPs, or apoplastic ROS, could contribute to Ca^2+^ influx in neighboring cells and assist wave transmission. Apoplastic ROS are implicated in the propagation of salt stress-induced rapid Ca^2+^ waves in roots and a ROS-assisted CICR model has been proposed, which can mathematically recapitulate the fast and constant wave speed (*14*). Additional mathematical modeling and experimental validation will help address these alternative hypotheses. Finally, our CICR model is based on the assumption that plasmodesmata (PD) do not restrict the diffusion of Ca^2+^ ions between cells (*14*). Cell-to-cell connectivity through PD is likely an important factor in determining the spatial pattern of defense responses (*63*). This is because PD-mediated cell-to-cell communication is downregulated by MAMPs such as chitin and flg22 and coordinated through unique PRR signaling complexes during PTI (*63*) as well as a target for bacterial type III secretion system effectors (*64*). Further studies are required to elucidate the role of PD in shaping the local traveling wave pattern and to explore whether heterogeneity in PD density and/or permeability (*63, 65*) influences the number and location of initiation sites.

Our work revealed that MAMP-induced local Ca^2+^ traveling waves emanate from a limited number of initiation sites or cells in cotyledon epidermal tissue. Similar patterns have been described in *Drosophila* wing discs during development-related Ca^2+^ wave propagation (*37*), and mimic the “leader-follower” model for collective chemosensing or information processing in multicellular models for animal systems (*38, 66*). The initiator or “leader” cells are defined as “first responders” to a signal, often displaying an above-average activity in generating Ca^2+^ spikes or oscillations (*37*) and/or having more connections or communications with neighboring cells (*38*). In our studies, it seems likely that initiator cells are more competent or robust for either MAMP perception or the activation of the Ca^2+^ encoding machinery. The observation that these cells were not more active during the run-in period for imaging prior to MAMP treatment suggests that the initiator cell behavior is more likely related to MAMP signal perception. This could be because they deploy a higher density of PRR receptors on their surface or, as our findings suggest, they have an average size that is larger compared to neighboring epidermal pavement cells. Single-cell heterogeneous responses to pathogen signals have been reported in other studies in Arabidopsis epidermal pavement, guard and root cells, using Ca^2+^ signatures as readouts (*26–28*), as well as in single-cell profiling studies with transcript signatures as readouts (*67, 68*). Recently, single-cell and spatial omics studies reveal a new cell population in plant leaves that serves as early responders to bacterial pathogen invasion with distinct transcript signatures compared to neighboring cells (*69*). Further research is required to understand the integration and communication of immune signals at the multicellular level in local plant tissues, utilizing space- and time-resolved technologies.

Another important feature we observed for MAMP-induced Ca^2+^ waves is their limited travel distance, typically reaching only 3–4 cells away from the initiation site. Given that Ca^2+^ signals are key players upstream of numerous signaling cascades involved in plant defense (*17, 20*), this limited spread likely represents a mechanism by which plants control and spatially restrict the local immune response during pathogen invasion. Research on both PTI and effector-triggered immunity (ETI) responses indicates that plant immune responses are tightly confined to only a few cell layers surrounding the pathogen invasion site. Zhu et al. (*68*) showed that upon infection of Arabidopsis rosette leaves with the bacterial pathogen *Pseudomonas syringae* pv. *tomato (Pst*), the expression of PTI-induced immune marker genes was predominantly limited to a few cells surrounding the local site of bacterial colonization. Plant defense responses related to ETI can also be spatially organized or confined, known as localized acquired resistance, and occur in a 1-to 2-mm zone surrounding a pathogen infection or treatment site when examined in tobacco leaves (*70, 71*). Recent studies report similar localized acquired responses where the strong induction of immune gene expression is only detected at the border of the infection area when Arabidopsis rosette leaves are infected with *Pst* (*72*). It has been proposed that the components of PTI signaling, such as calcium influxes, apoplastic ROS production, or DAMP release, likely contribute to the spatial pattern in local acquired resistance (*72*). Similarly, spatial restriction of immune responses has been reported in plant roots through controlling the expression pattern of PRR receptors in different tissue and cell types (*73–75*). This spatial confinement of pathogen-induced defense responses likely reflects a strategy by plants to strike a balance between growth and defense (*72, 74*). Minimizing the inhibitory effects of pathogen attack on plant growth is critical, given that plant organs, such as roots and shoots, are continuously growing throughout the plant’s life cycle. If defense responses are too widespread or prolonged, they could significantly impair plant growth and development, leading to reductions in overall biomass or crop yield. It is therefore important to further investigate the cellular and molecular mechanisms underlying the generation and attenuation of the PTI-induced local traveling waves, leveraging high spatiotemporal resolution Ca^2+^ imaging and computational modeling approaches, as demonstrated in this study.

## MATERIALS AND METHODS

### Plant materials and growth conditions

All *Arabidopsis thaliana* transgenic or mutant lines used in this study were in Col-0 ecotype. The *Arabidopsis thaliana* R-GECO1 reporter line (*27*) was kindly provided by Melanie Krebs (University of Heidelberg). Homozygous *fls2* and wild-type siblings expressing R-GECO1 were obtained by crossing the *fls2* mutant (SALK_062054) with R-GECO1 and recovery from an F2 population. Selfed F3 lines were used for all experiments. Seeds were surface sterilized and stratified at 4°C for 2 to 3 days on half-strength Murashige and Skoog medium supplemented with 1% (w/v) sucrose and 1% (w/v) agar (pH 5.8). Plants were grown under a light intensity of 120−140 μmol m^−2^ s^−1^ provided by Philips F32T8/L941 Deluxe Cool White bulbs, under long-day conditions (16 h light/8 h dark) at 21°C. Seven-day-old seedlings were used for calcium imaging.

### Chemicals and elicitors

The MAMP/DAMP peptides, flg22 (RP19986, GenScript, Piscataway, NJ, USA) and Pep1 (NeoBioSci, Cambridge, MA, USA), were dissolved in 1 X phosphate-buffered saline (PBS) buffer to make 1 mM stock solutions and stored at -80°C. Stock solutions of 10 mg/mL chitin were prepared by suspending chitin powder from shrimp shells (C9752, Sigma-Aldrich, St. Louis, MO, USA) in 1 X PBS buffer, followed by sonication for 3 × 30 s. The suspension was then centrifuged and the supernatant aliquoted and stored at -80°C. Stock solutions of 1 M LaCl_3_ (203521, Sigma-Aldrich) and 25 mM BAPTA-AM (A1076, Sigma-Aldrich) were made in water and DMSO, respectively. For glutamate stock solutions, L-glutamic acid (Thermo Fisher Scientific, Waltham, MA, USA) was dissolved in 0.05% (w/v) MES (pH adjusted to 5.3 using KOH) to make a 50 mM stock solution.

### Calcium imaging

For live-cell imaging experiments, epidermal pavement cells on the abaxial side of Arabidopsis cotyledons and from the central to apical region of the cotyledon epidermis (fig. S1E) were used exclusively. Seven-day-old cotyledons were detached from seedlings grown on plates one day prior to imaging and floated in sterile water overnight under light to minimize wound-induced responses. Cotyledons were then mounted in 2-well chambered coverslips (ibidi GmbH, Gräfelfing, Germany) for bottom imaging, with a chamber design based on Keinath et al. (*27*) with modifications (fig. S1A). Cotyledons were soaked in water in the chamber and a cotton pad was placed on top as a spacer. A thin agar block with a central hole was placed on top of the cotton pad to stabilize the sample without applying direct physical pressure to the cotyledon. The assembled chamber was allowed to rest in the microscope room for 40–60 min prior to imaging. Spinning-disk confocal microscopy (SDCM) with a Yokogawa scanner unit CSU-X1-A1 (Hamamatsu Photonics, Hamamatsu, Japan) mounted on an Olympus IX-83 microscope, equipped with a 10X 0.4–numerical aperture (NA) UPLXAPO objective (Olympus, Tokyo, Japan) and an Andor iXon Ultra 897BV EMCCD camera (Andor Technology, Belfast, UK), was used for all Ca^2+^ imaging experiments. R-GECO1 was excited with a 561-nm laser, and emission was collected through a 610/37-nm filter. Cotyledon epidermal cells were imaged at a single focal plane with 5-s acquisition intervals for a total of 40 min. Samples were first imaged for 10 min without any treatment as a run-in period for any laser light-induced Ca^2+^ activities. Samples with high spontaneous Ca^2+^ activity or that failed to maintain a normal background Ca^2+^ level during the first 10 min were discarded. For global MAMP/DAMP treatments, image acquisition was paused after the 120^th^ frame, and a two-fold concentration of elicitor solution diluted in water was gently added to the central hole of the imaging chamber in a 1:1 volume ratio. Image acquisition was then resumed for another 30 min. For inhibitor pre-treatments, a two-fold concentration of inhibitor solution diluted in water was added to the imaging chamber in a 1:1 volume ratio at desired time points prior to imaging.

To image wound-induced Ca^2+^ signals, cotyledons were prepared, mounted in imaging chambers, and imaged with the same microscopy settings as described above. An Andor Mosaic 3 photomanipulation module (Andor Technology) integrated into the SDCM system was used for single-cell laser ablation and wound stimulation. A random cell in the cotyledon epidermis was irradiated with a high-power (1.3 W) 445-nm laser at 80% power for 1 s with an ROI size of 16 × 16 pixels. Time-lapse image series were acquired at 2-s intervals for 7 min.

### Image analysis

Image processing and extraction of Ca^2+^ intensity traces were performed in FIJI (*76*). An average intensity projection generated from individual timelapse image series was used for individual cell segmentation. The puzzle piece-shaped epidermal cells were segmented by either manual tracing of the cell borders with the freehand line tool in FIJI or using the deep-learning based software LeafNet (*34*). For LeafNet segmentation, the Stained Denoiser and LeafSeg methods were used. The annotated output image from LeafNet was converted into a binary image and skeletonized in FIJI, and any segmentation errors were manually corrected in FIJI. The borders of individual cells were selected automatically with the Wand tracing tool and saved as ROIs in the ROI Manager (fig. S1, B and C). The ROIs were selections of the whole cell area and were used for measuring cell areas. For extracting calcium intensity profiles, the area selections were converted into line selections with a width of 2 pixels, so that the Ca^2+^ intensity was only extracted from the borders of each cell representing the thin cytoplasmic layer in the cell periphery. The Plot Z-axis Profile function in FIJI was used to extract the average Ca^2+^ intensity traces from the ROIs in the timelapse image series. The raw fluorescence values were converted into the ratio ΔF/F_0_ or (F – F_0_)/F_0_ after subtracting the background. The F_0_ represents the baseline fluorescence calculated from the average intensity of a stable period in the first 10–12 min of the imaging period prior to elicitor treatment.

Peak analysis of Ca^2+^ traces was performed in MATLAB with the *findpeaks* function (code link: https://github.com/SADDLab/Calcium_manuscript_2024_Image_Analysis_Code.git). The minimum peak height and prominence was set to 0.5, and the minimum peak width was set to 40 for peak identification. The mean peak interval was calculated as the mean interval between peaks in a trace with the meanCycle argument in MATLAB. The total response time was calculated as the duration from the 1^st^ peak to the last peak using peak width intercepts, and the area under the curve (AUC) was calculated accordingly as the area within the total response time period (fig. S1D).

For analysis of intercellular Ca^2+^ wave speed, a FIJI macro was developed based on Bellandi et al. (*10*) to scan a wave using a series of concentric circles instead of a series of radial line scans (code link: https://github.com/SADDLab/Calcium_manuscript_2024_Image_Analysis_Code.git).

Specifically, ROIs of the individual waves were cropped from the original video and a set of concentric circles was generated from the center of a wave, with the radius increased by 5 pixels at each step to cover the whole wave travel range (fig. S7A). The mean fluorescence intensity was extracted from each circular ROI at each time point (fig. S7B). The fluorescence traces were smoothed in MATLAB with the “loess” method, and a single peak was detected. The peak value corresponded to the time a wave travels a certain distance from the origin (radius of the circle). The distance and peak values (time) were plotted, and a linear curve was fitted to calculate the wave speed (fig. S7C). Only curves that could be fitted with a linear function with R^2^ > 0.9 were used for speed analyses. For laser ablation-induced wave speed analysis, images were analyzed using the same method as described above, except that the distance vs. time curve was fitted with a polynomial of degree 3, and the derivatives of the polynomial were calculated in MATLAB and used as wave speed (Fig. 3, C and D).

### Statistical analysis

Measurements of multiple cells or waves from a single cotyledon were averaged as one data point representing a single biological sample, and at least 3 samples were analyzed per treatment or per biological repeat. Data from 3 independent experiments were binned together for statistical analysis. One-way ANOVA and Tukey’s HSD post hoc tests were performed in RStudio to determine significance among different treatments.

### Computational modeling of calcium-induced calcium release (CICR)

We utilized the calcium-induced calcium-release mathematical model proposed by

Dawson et. al (*41*),

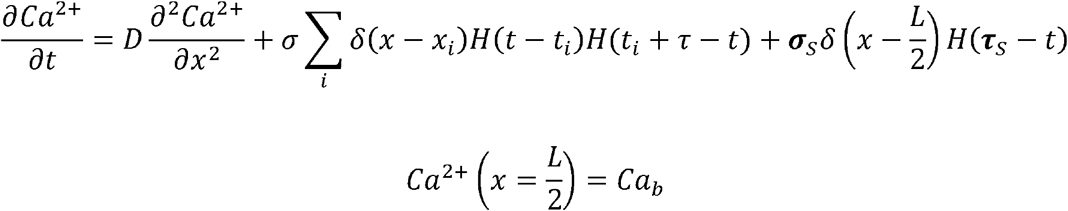

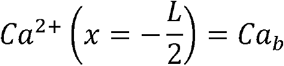

In the equations, *t*_*i*_denotes the time at which calcium (Ca^2+^, in μM) channel becomes activated in response to local Ca^2+^ levels. The model is solved in 1D and the position of Ca^2+^ channel along the x-axis is denoted by x_i_ (in μm). When the local Ca^2+^concentration exceeds a threshold value of *Ca*_*T*_ (in μM), the Ca^2+^ channel is activated with a refractory time of τminutes. The strength of the Ca^2+^ release is represented by the rate at which Ca^2+^ is released upon the opening, denoted as σ. δ and *H* denotes Dirac-Delta and Heaviside functions respectively. *σ*_*s*_ represents the flux at the center of domain which is used to represent the initiator site or source *τ*_*s*_ represents the time for which the point source is active (Table S1).

The model was solved in Python using an explicit finite difference method (*77*). The total domain length (*L*) was 700 µm, with a spatial resolution (*dx*) of 0.2 μm. The time step (*dt*) used for the finite difference Euler method was 10^−5^ minutes. The Ca^2+^ channels are spatially patterned and separated by *d* μm. To manage boundary conditions, the CNGC channels are distributed only within the domain length of -250 µm to 250 µm. Beyond this range, calcium spreads solely by free diffusion to prevent calcium accumulation at the boundaries (fig. S9).

The modeling of a Ca^2+^-dependent sink assumes that Ca^2+^ binding occurs rapidly, keeping the process in equilibrium. In this model, we consider that each binding event requires *n* Ca^2+^ ions, with forward and backward rate constants *k*_*f*_ and *k*_*b*_, respectively, for Ca^2+^ binding and unbinding. Based on these assumptions, we derive

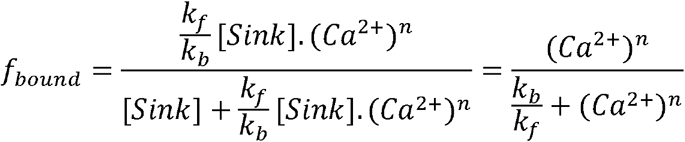

where *f* _*bound*_ represents the fraction of sink type proteins that are Ca^2+^ bound, active and sequestering Ca^2+^ out of cytoplasm. Using the above we define the flux through sink as,

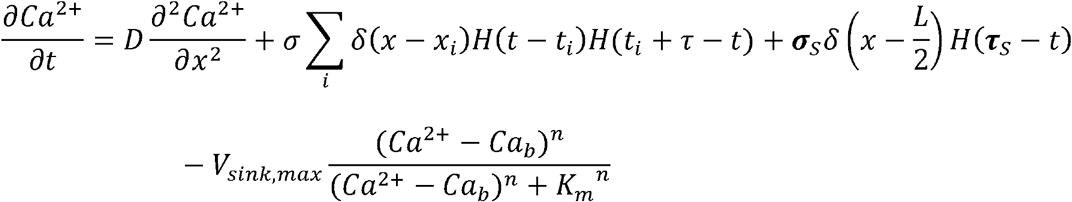

where *v*_*sink,max*_ represents the maximum flux through sink. A summary of known model parameters from different organisms with distinct traveling waves and their relevance can be found in Table S1. Code for the modeling work can be found in link: https://github.com/SADDLab/Calcium_manuscript_2024_1D_wave_model.git.

## Supplementary Materials

Fig. S1. Pipeline for calcium imaging and single-cell calcium signature analysis.

Fig. S2. Pre-treatment with LaCl_3_ or BAPTA-AM abolishes flg22-induced cytosolic Ca^2+^ oscillations.

Fig. S3. The flg22-induced cytosolic Ca^2+^ oscillations are abolished in *fls2*.

Fig. S4. Initiator cells are randomly distributed in epidermal tissue and are larger than neighboring cells.

Fig. S5. Representative traces of Ca^2+^ intensity ratio (ΔF/F_0_) for initiator (N) and neighboring (N+1 and N+2) cells following global treatment with 1 µM flg22.

Fig. S6. Calcium signatures and wave propagation patterns in cotyledon epidermal cells treated with various MAMPs or DAMPs.

Fig. S7. A method for analysis of the speed of intercellular Ca^2+^ traveling waves in the cotyledon epidermis.

Fig. S8. Single-cell Ca^2+^ signatures in response to global glutamate treatment differ from those elicited by MAMPs.

Fig. S9. Mathematical modeling of 1D wave propagation through Ca^2+^-induced Ca^2+^ release.

Fig. S10. Local sensitivity and correlation analysis of model parameters. Fig. S11. Extended analysis for mathematical model with a sink.

Table S1. List of CICR model parameters.

Movie S1. Cytosolic Ca^2+^ dynamics in cotyledon epidermal cells following global treatment with flg22.

Movie S2. Local traveling waves of Ca^2+^ induced by flg22 are initiated from a small subset of cells randomly distributed in epidermal tissue.

Movie S3. Cytosolic Ca^2+^ dynamics in cotyledon epidermal cells following global treatment with chitin.

Movie S4. Cytosolic Ca^2+^ dynamics in cotyledon epidermal cells following global treatment with Pep1.

Movie S5. Wounding of a single epidermal cell with laser irradiation induces a Ca^2+^ traveling wave.

Movie S6. Calcium traveling waves in cotyledon epidermal cells elicited by 10 nM flg22. Supplementary movie legends.

## Acknowledgments

We thank Melanie Krebs (University of Heidelberg) for sharing the R-GECO1 transgenic line and Sheng Luan (University of California, Berkeley) for providing the *cngc2/4* double mutant. The authors are grateful to Hongbing Luo (Purdue) for excellent care and maintenance of plant materials.

## Author contributions

Conceptualization: CJS, EP, DMU, AIP

Funding acquisition: DMU, CJS

Methodology: CJS, EP, WZ, NK

Data acquisition and analysis: WZ, JRH, AH, NK

Computational modeling: NK, EP Visualization: WZ, NK

Writing – original draft: WZ, NK, CJS

Writing – review & editing: WZ, NK, CJS, EP, AIP, DMU

## Data and materials availability

Code for data analysis and computational remodeling are available in Github and links are provided in the Materials and Methods section.

## References and Notes

1. D. E. Clapham, Calcium signaling. Cell 131, 1047–1058 (2007).

2. S. Luan, C. Wang, Calcium signaling mechanisms across kingdoms. Annu. Rev. Cell Dev. Biol. 37, 311–340 (2021).

3. A. N. Dodd, J. Kudla, D. Sanders, The language of calcium signaling. Annu. Rev. Plant Biol. 61, 593–620 (2010).

4. C. Allan, R. J. Morris, C.-N. Meisrimler, Encoding, transmission, decoding, and specificity of calcium signals in plants. J. Exp. Bot. 73, 3372–3385 (2022).

5. K. H. Edel, E. Marchadier, C. Brownlee, J. Kudla, A. M. Hetherington, The evolution of calcium-based signalling in plants. Curr. Biol. 27, R667–R679 (2017).

6. J. Kudla, D. Becker, E. Grill, R. Hedrich, M. Hippler, U. Kummer, M. Parniske, T. Romeis, K. Schumacher, Advances and current challenges in calcium signaling. New Phytol. 218, 414–431 (2018).

7. W. Tian, C. Wang, Q. Gao, L. Li, S. Luan, Calcium spikes, waves and oscillations in plant development and biotic interactions. Nat. Plants 6, 750–759 (2020).

8. Y. Fichman, R. Mittler, Integration of electric, calcium, reactive oxygen species and hydraulic signals during rapid systemic signaling in plants. Plant J. 107, 7–20 (2021).

9. R. Hilleary, S. Gilroy, Systemic signaling in response to wounding and pathogens. Curr. Opin. Plant Biol. 43, 57–62 (2018).

10. A. Bellandi, D. Papp, A. Breakspear, J. Joyce, M. G. Johnston, J. de Keijzer, E. C. Raven, M. Ohtsu, T. R. Vincent, A. J. Miller, D. Sanders, S. A. Hogenhout, R. J. Morris, C. Faulkner, Diffusion and bulk flow of amino acids mediate calcium waves in plants. Sci. Adv. 8, eabo6693 (2022).

11. J. Guo, J. He, K. Dehesh, X. Cui, Z. Yang, CamelliA-based simultaneous imaging of Ca^2+^ dynamics in subcellular compartments. Plant Physiol. 188, 2253–2271 (2022).

12. M. Matsumura, M. Nomoto, T. Itaya, Y. Aratani, M. Iwamoto, T. Matsuura, Y. Hayashi, T. Mori, M. J. Skelly, Y. Y. Yamamoto, T. Kinoshita, I. C. Mori, T. Suzuki, S. Betsuyaku, S.H. Spoel, M. Toyota, Y. Tada, Mechanosensory trichome cells evoke a mechanical stimuli– induced immune response in Arabidopsis thaliana. Nat. Commun. 13, 1216 (2022).

13. A. H. Howell, C. Völkner, P. McGreevy, K. H. Jensen, R. Waadt, S. Gilroy, H.-H. Kunz, W. S. Peters, M. Knoblauch, Pavement cells distinguish touch from letting go. Nat. Plants 9, 877–882 (2023).

14. M. J. Evans, W.-G. Choi, S. Gilroy, R. J. Morris, A ROS-assisted calcium wave dependent on the AtRBOHD NADPH oxidase and TPC1 cation channel propagates the systemic response to salt stress. Plant Physiol. 171, 1771–1784 (2016).

15. Y. Fichman, R. J. Myers, D. G. Grant, R. Mittler, Plasmodesmata-localized proteins and ROS orchestrate light-induced rapid systemic signaling in Arabidopsis. Sci. Signal. 14, eabf0322 (2021).

16. M. Toyota, D. Spencer, S. Sawai-Toyota, W. Jiaqi, T. Zhang, A. J. Koo, G. A. Howe, S. Gilroy, Glutamate triggers long-distance, calcium-based plant defense signaling. Science 361, 1112–1115 (2018).

17. H. Seybold, F. Trempel, S. Ranf, D. Scheel, T. Romeis, J. Lee, Ca^2+^ signalling in plant immune response: from pattern recognition receptors to Ca^2+^ decoding mechanisms. New Phytol. 204, 782–790 (2014).

18. I. Albert, C. Hua, T. Nürnberger, R. N. Pruitt, L. Zhang, Surface sensor systems in plant immunity. Plant Physiol. 182, 1582–1596 (2020).

19. T. A. DeFalco, C. Zipfel, Molecular mechanisms of early plant pattern-triggered immune signaling. Mol. Cell 81, 3449–3467 (2021).

20. Y. Jiang, P. Ding, Calcium signaling in plant immunity: a spatiotemporally controlled symphony. Trends Plant Sci. 28, 74–89 (2023).

21. P. Köster, T. A. DeFalco, C. Zipfel, Ca^2+^ signals in plant immunity. EMBO J. 41, e110741 (2022).

22. P. Yuan, E. Jauregui, L. Du, K. Tanaka, B. Poovaiah, Calcium signatures and signaling events orchestrate plant–microbe interactions. Curr. Opin. Plant Biol. 38, 173–183 (2017).

23. D. Aldon, M. Mbengue, C. Mazars, J.-P. Galaud, Calcium signalling in plant biotic interactions. Int. J. Mol. Sci. 19, 665 (2018).

24. S. Ranf, L. Eschen-Lippold, P. Pecher, J. Lee, D. Scheel, Interplay between calcium signalling and early signalling elements during defence responses to microbe-or damage-associated molecular patterns. Plant J. 68, 100–113 (2011).

25. M. Grenzi, F. Resentini, S. Vanneste, M. Zottini, A. Bassi, A. Costa, Illuminating the hidden world of calcium ions in plants with a universe of indicators. Plant Physiol. 187, 550–571 (2021).

26. B. Eichstädt, S. Lederer, F. Trempel, X. Jiang, T. Guerra, R. Waadt, J. Lee, A. Liese, T. Romeis, Plant immune memory in systemic tissue does not involve changes in rapid calcium signaling. Front. Plant Sci. 12 (2021).

27. N. F. Keinath, R. Waadt, R. Brugman, J. I. Schroeder, G. Grossmann, K. Schumacher, M. Krebs, Live cell imaging with R-GECO1 sheds light on flg22- and chitin-induced transient [Ca^2+^]cyt patterns in Arabidopsis. Mol. Plant 8, 1188–1200 (2015).

28. K. Thor, E. Peiter, Cytosolic calcium signals elicited by the pathogen-associated molecular pattern flg22 in stomatal guard cells are of an oscillatory nature. New Phytol. 204, 873–881 (2014).

29. M. Beck, J. Zhou, C. Faulkner, D. MacLean, S. Robatzek, Spatio-temporal cellular dynamics of the Arabidopsis flagellin receptor reveal activation status-dependent endosomal sorting. Plant Cell 24, 4205–4219 (2012).

30. L. Cao, W. Wang, W. Zhang, C. J. Staiger, Lipid signaling requires ROS production to elicit actin cytoskeleton remodeling during plant innate immunity. Int. J. Mol. Sci. 23, 2447 (2022).

31. R. Hilleary, J. Paez-Valencia, C. Vens, M. Toyota, M. Palmgren, S. Gilroy, Tonoplast-localized Ca^2+^ pumps regulate Ca^2+^ signals during pattern-triggered immunity in Arabidopsis thaliana. Proc. Natl. Acad. Sci. 117, 18849–18857 (2020).

32. G. Felix, J. D. Duran, S. Volko, T. Boller, Plants have a sensitive perception system for the most conserved domain of bacterial flagellin. Plant J. 18, 265–276 (1999).

33. L. Gómez-Gómez, T. Boller, FLS2: an LRR receptor–like kinase involved in the perception of the bacterial elicitor flagellin in Arabidopsis. Mol. Cell 5, 1003–1011 (2000).

34. S. Li, L. Li, W. Fan, S. Ma, C. Zhang, J. C. Kim, K. Wang, E. Russinova, Y. Zhu, Y. Zhou, LeafNet: a tool for segmenting and quantifying stomata and pavement cells. Plant Cell 34, 1171–1188 (2022).

35. N. Shibuya, E. Minami, Oligosaccharide signalling for defence responses in plant. Physiol. Mol. Plant Pathol. 59, 223–233 (2001).

36. A. Huffaker, G. Pearce, C. A. Ryan, An endogenous peptide signal in Arabidopsis activates components of the innate immune response. Proc. Natl. Acad. Sci. 103, 10098–10103 (2006).

37. D. K. Soundarrajan, F. J. Huizar, R. Paravitorghabeh, T. Robinett, J. J. Zartman, From spikes to intercellular waves: Tuning intercellular calcium signaling dynamics modulates organ size control. PLOS Comput. Biol. 17, e1009543 (2021).

38. V. Salem, L. D. Silva, K. Suba, E. Georgiadou, S. Neda Mousavy Gharavy, N. Akhtar, A. Martin-Alonso, D. C. A. Gaboriau, S. M. Rothery, T. Stylianides, G. Carrat, T. J. Pullen, S.P. Singh, D. J. Hodson, I. Leclerc, A. M. J. Shapiro, P. Marchetti, L. J. B. Briant, W. Distaso, N. Ninov, G. A. Rutter, Leader β-cells coordinate Ca^2+^ dynamics across pancreatic islets in vivo. Nat. Metab. 1, 615–629 (2019).

39. L. F. Jaffe, Classes and mechanisms of calcium waves. Cell Calcium 14, 736–745 (1993).

40. L. F. Jaffe, On the conservation of fast calcium wave speeds. Cell Calcium 32, 217–229 (2002).

41. S. P. Dawson, J. Keizer, J. E. Pearson, Fire–diffuse–fire model of dynamics of intracellular calcium waves. Proc. Natl. Acad. Sci. 96, 6060–6063 (1999).

42. M. D. Mckay, R. J. Beckman, W. J. Conover, A comparison of three methods for selecting values of input variables in the analysis of output from a computer code. Technometrics 42, 55–61 (2000).

43. N. Frei dit Frey, M. Mbengue, M. Kwaaitaal, L. Nitsch, D. Altenbach, H. Häweker, R. Lozano-Duran, M. F. Njo, T. Beeckman, B. Huettel, J. W. Borst, R. Panstruga, S. Robatzek, Plasma membrane calcium ATPases are important components of receptor-mediated signaling in plant immune responses and development. Plant Physiol. 159, 798– 809 (2012).

44. Z. Li, J. F. Harper, C. Weigand, J. Hua, Resting cytosol Ca^2+^ level maintained by Ca^2+^ pumps affects environmental responses in Arabidopsis. Plant Physiol. 191, 2534–2550 (2023).

45. A. Politi, L. D. Gaspers, A. P. Thomas, T. Höfer, Models of IP3 and Ca^2+^ oscillations: Frequency encoding and identification of underlying feedbacks. Biophys. J. 90, 3120–3133 (2006).

46. W. Tian, C. Hou, Z. Ren, C. Wang, F. Zhao, D. Dahlbeck, S. Hu, L. Zhang, Q. Niu, L. Li, B. J. Staskawicz, S. Luan, A calmodulin-gated calcium channel links pathogen patterns to plant immunity. Nature 572, 131–135 (2019).

47. M. K. Meena, R. Prajapati, D. Krishna, K. Divakaran, Y. Pandey, M. Reichelt, M. K. Mathew, W. Boland, A. Mithöfer, J. Vadassery, The Ca^2+^ channel CNGC19 regulates arabidopsis defense against spodoptera herbivory. Plant Cell 31, 1539–1562 (2019).

48. S. Johns, T. Hagihara, M. Toyota, S. Gilroy, The fast and the furious: rapid long-range signaling in plants. Plant Physiol. 185, 694–706 (2021).

49. T. Hander, Á. D. Fernández-Fernández, R. P. Kumpf, P. Willems, H. Schatowitz, D. Rombaut, A. Staes, J. Nolf, R. Pottie, P. Yao, A. Gonçalves, B. Pavie, T. Boller, K. Gevaert, F. Van Breusegem, S. Bartels, S. Stael, Damage on plants activates Ca^2+^-dependent metacaspases for release of immunomodulatory peptides. Science 363, eaar7486 (2019).

50. L. Bacete, H. Mélida, E. Miedes, A. Molina, Plant cell wall-mediated immunity: cell wall changes trigger disease resistance responses. Plant J. 93, 614–636 (2018).

51. E. Peiter, F. J. M. Maathuis, L. N. Mills, H. Knight, J. Pelloux, A. M. Hetherington, D. Sanders, The vacuolar Ca^2+^-activated channel TPC1 regulates germination and stomatal movement. Nature 434, 404–408 (2005).

52. W.-G. Choi, M. Toyota, S.-H. Kim, R. Hilleary, S. Gilroy, Salt stress-induced Ca^2+^ waves are associated with rapid, long-distance root-to-shoot signaling in plants. Proc. Natl. Acad. Sci. 111, 6497–6502 (2014).

53. S. Ranf, P. Wünnenberg, J. Lee, D. Becker, M. Dunkel, R. Hedrich, D. Scheel, P. Dietrich, Loss of the vacuolar cation channel, AtTPC1, does not impair Ca^2+^ signals induced by abiotic and biotic stresses. Plant J. 53, 287–299 (2008).

54. Y. Ma, R. K. Walker, Y. Zhao, G. A. Berkowitz, Linking ligand perception by PEPR pattern recognition receptors to cytosolic Ca^2+^ elevation and downstream immune signaling in plants. Proc. Natl. Acad. Sci. 109, 19852–19857 (2012).

55. Y. Ma, K. Garrido, R. Ali, G. A. Berkowitz, Phenotypes of cyclic nucleotide-gated cation channel mutants: probing the nature of native channels. Plant J. 113, 1223–1236 (2023).

56. P. Dietrich, W. Moeder, K. Yoshioka, Plant cyclic nucleotide-gated channels: new insights on their functions and regulation. Plant Physiol. 184, 27–38 (2020).

57. M. Bjornson, P. Pimprikar, T. Nürnberger, C. Zipfel, The transcriptional landscape of Arabidopsis thaliana pattern-triggered immunity. Nat. Plants 7, 579–586 (2021).

58. C. Espinoza, Y. Liang, G. Stacey, Chitin receptor CERK1 links salt stress and chitin-triggered innate immunity in Arabidopsis. Plant J. 89, 984–995 (2017).

59. K. Thor, S. Jiang, E. Michard, J. George, S. Scherzer, S. Huang, J. Dindas, P. Derbyshire, N. Leitão, T. A. DeFalco, P. Köster, K. Hunter, S. Kimura, J. Gronnier, L. Stransfeld, Y. Kadota, C. A. Bücherl, M. Charpentier, M. Wrzaczek, D. MacLean, G. E. D. Oldroyd, F. L. H. Menke, M. R. G. Roelfsema, R. Hedrich, J. Feijó, C. Zipfel, The calcium-permeable channel OSCA1.3 regulates plant stomatal immunity. Nature 585, 569–573 (2020).

60. C.-Y. Hung, P. Aspesi Jr, M. R. Hunter, A. W. Lomax, I. Y. Perera, Phosphoinositide-signaling is one component of a robust plant defense response. Front. Plant Sci. 5 (2014).

61. A. M. Murphy, B. Otto, C. A. Brearley, J. P. Carr, D. E. Hanke, A role for inositol hexakisphosphate in the maintenance of basal resistance to plant pathogens. Plant J. 56, 638–652 (2008).

62. Y. Ma, Y. Zhao, G. A. Berkowitz, Intracellular Ca^2+^ is important for flagellin-triggered defense in Arabidopsis and involves inositol polyphosphate signaling. J. Exp. Bot. 68, 3617–3628 (2017).

63. E. E. Tee, C. Faulkner, Plasmodesmata and intercellular molecular traffic control. New Phytol. 243, 32–47 (2024).

64. K. Aung, P. Kim, Z. Li, A. Joe, B. Kvitko, J. R. Alfano, S. Y. He, Pathogenic bacteria target plant plasmodesmata to colonize and invade surrounding tissues. Plant Cell 32, 595– 611 (2020).

65. C. Gao, X. Liu, N. De Storme, K. H. Jensen, Q. Xu, J. Yang, X. Liu, S. Chen, H. J. Martens, A. Schulz, J. Liesche, Directionality of plasmodesmata-mediated transport in arabidopsis leaves supports auxin channeling. Curr. Biol. 30, 1970-1977.e4 (2020).

66. G. Li, R. LeFebre, A. Starman, P. Chappell, A. Mugler, B. Sun, Temporal signals drive the emergence of multicellular information networks. Proc. Natl. Acad. Sci. U. S. A. 119, e2202204119 (2022).

67. B. Tang, L. Feng, M. T. Hulin, P. Ding, W. Ma, Cell-type-specific responses to fungal infection in plants revealed by single-cell transcriptomics. Cell Host Microbe 31, 1732-1747.e5 (2023).

68. J. Zhu, S. Lolle, A. Tang, B. Guel, B. Kvitko, B. Cole, G. Coaker, Single-cell profiling of Arabidopsis leaves to Pseudomonas syringae infection. Cell Rep. 42, 112676 (2023).

69. T. Nobori, A. Monell, T. A. Lee, Y. Sakata, S. Shirahama, J. Zhou, J. R. Nery, A. Mine, J. R. Ecker, A rare PRIMER cell state in plant immunity. Nature, 1–9 (2025).

70. S. Dorey, F. Baillieul, M.-A. Pierrel, P. Saindrenan, B. Fritig, S. Kauffmann, Spatial and temporal induction of cell death, defense genes, and accumulation of salicylic acid in tobacco leaves reacting hypersensitively to a fungal glycoprotein elicitor. Mol. Plant Microbe Interact. 10, 646–655 (1997).

71. A. F. Ross, Localized acquired resistance to plant virus infection in hypersensitive hosts. Virology 14, 329–339 (1961).

72. P. Jacob, J. Hige, J. L. Dangl, Is localized acquired resistance the mechanism for effector-triggered disease resistance in plants? Nat. Plants 9, 1184–1190 (2023).

73. M. Beck, I. Wyrsch, J. Strutt, R. Wimalasekera, A. Webb, T. Boller, S. Robatzek, Expression patterns of FLAGELLIN SENSING 2 map to bacterial entry sites in plant shoots and roots. J. Exp. Bot. 65, 6487–6498 (2014).

74. A. Emonet, F. Zhou, J. Vacheron, C. M. Heiman, V. Dénervaud Tendon, K.-W. Ma, P. Schulze-Lefert, C. Keel, N. Geldner, Spatially restricted immune responses are required for maintaining root meristematic activity upon detection of bacteria. Curr. Biol. 31, 1012-1028.e7 (2021).

75. F. Zhou, A. Emonet, V. D. Tendon, P. Marhavy, D. Wu, T. Lahaye, N. Geldner, Co-incidence of damage and microbial patterns controls localized immune responses in roots. Cell 180, 440-453.e18 (2020).

76. J. Schindelin, I. Arganda-Carreras, E. Frise, V. Kaynig, M. Longair, T. Pietzsch, S. Preibisch, C. Rueden, S. Saalfeld, B. Schmid, J.-Y. Tinevez, D. J. White, V. Hartenstein, K. Eliceiri, P. Tomancak, A. Cardona, Fiji: an open-source platform for biological-image analysis. Nat. Methods 9, 676–682 (2012).

77. Smith, Gordon D., Numerical Solution of Partial Differential Equations: Finite Difference Methods. (Oxford University Press, 1985).

