## Supplemental Figures for "Local traveling waves of cytosolic calcium elicited by defense signals or wounding are propagated by distinct mechanisms"

### Supplementary Materials

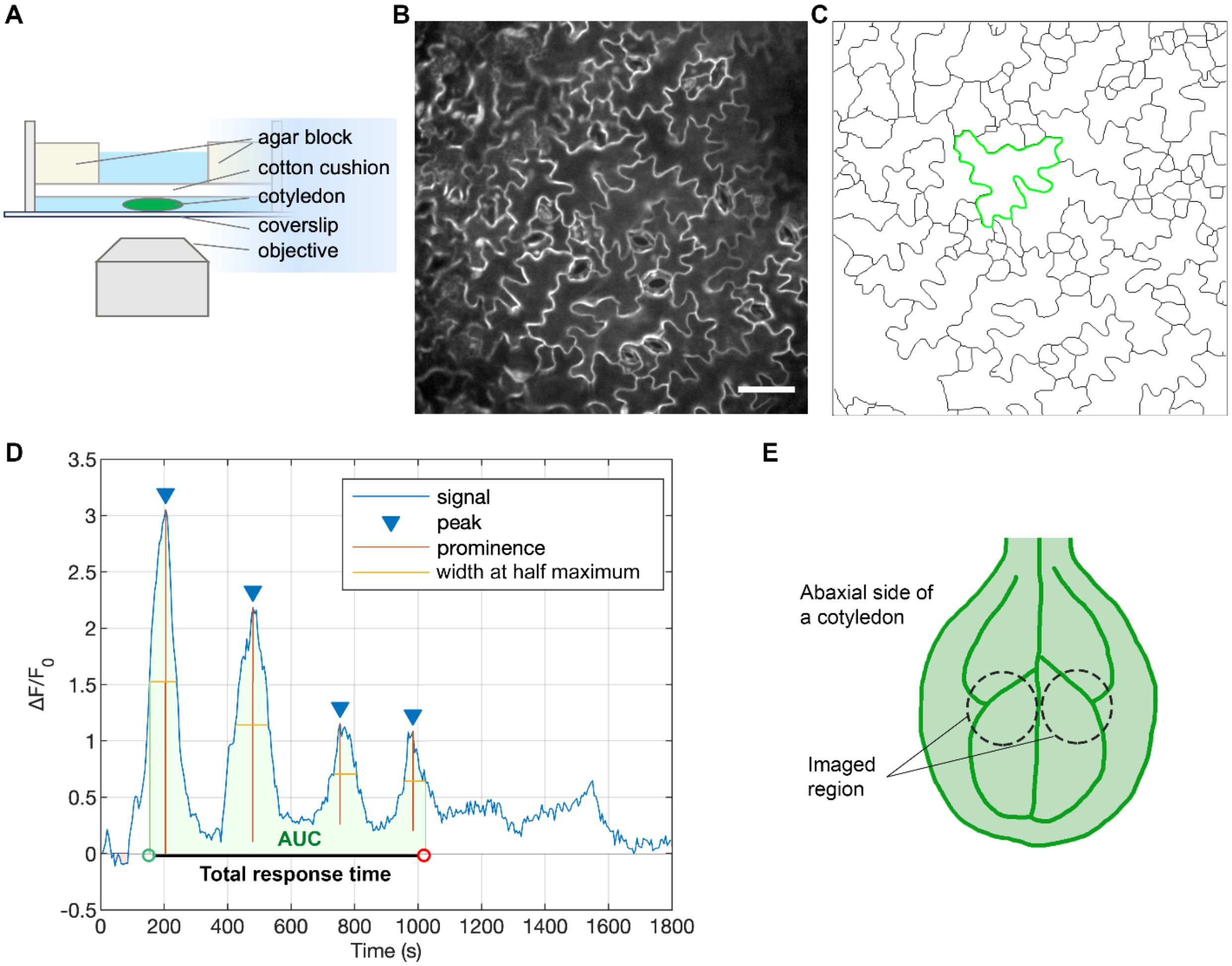


**Fig. S1. Pipeline for calcium imaging and single-cell calcium signature analysis.** (**A**) The design of a chamber for live-cell imaging of excised Arabidopsis cotyledons by spinning disk confocal microscopy (SDCM) with an inverted objective is diagrammed. (**B**) A representative image of Arabidopsis cotyledon epidermal cells expressing the genetically-encoded calcium reporter, R-GECO1 (Keinath et al., 2015). Image shown is an average intensity projection of a 30-min timelapse series collected with SDCM. Bar = 50 µm. (**C**) Segmented image of (B) using LeafNet (Li et al., 2022) and single-cell perimeter extraction (green line) as the ROI for calcium fluorescent intensity measurement. (**D**) MAMP-induced calcium intensity trace from a single cell, extracted as shown in (C), was converted into $\Delta F/F_0$ and the calcium peak features analyzed with the MATLAB *findpeaks* function. AUC: area under the curve. (**E**) A diagram shows the regions on the abaxial face of a 7-d-old cotyledon that were used for $Ca^{2+}$ imaging.

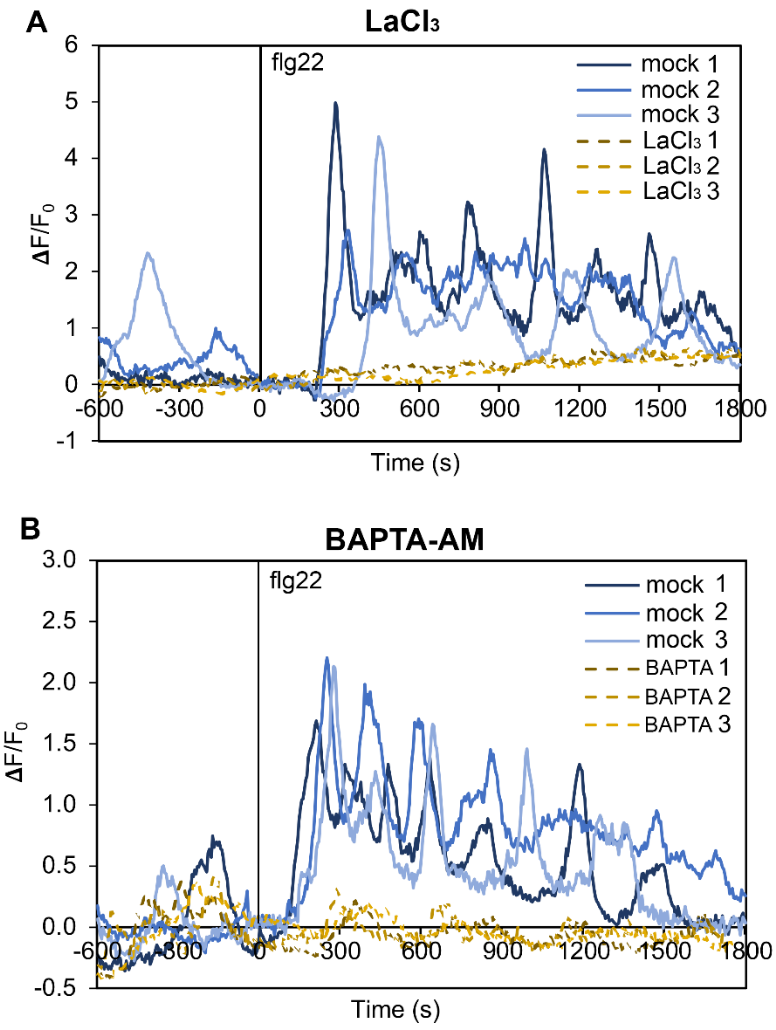


**Fig. S2. Pre-treatment with $LaCl_3$ or BAPTA-AM abolishes flg22-induced cytosolic $Ca^{2+}$ oscillations.** (**A**) Arabidopsis cotyledons expressing R-GECO1 were pre-treated with mock or 500 µM $LaCl_3$ for 30 min followed by 1 µM flg22 treatment. Fluorescence intensity ratio profiles were extracted from individual cells and blue lines represent mock-treated cells, whereas brown dashed lines represent $LaCl_3$-treated cells. (**B**) Cotyledons were pre-treated with mock or 25 µM BAPTA-AM for 30 min followed by 1 µM flg22 treatment. Blue lines represent mock-treated cells and brown dashed lines represent BAPTA-AM-treated cells. Data shown are from 3 cotyledons per treatment.

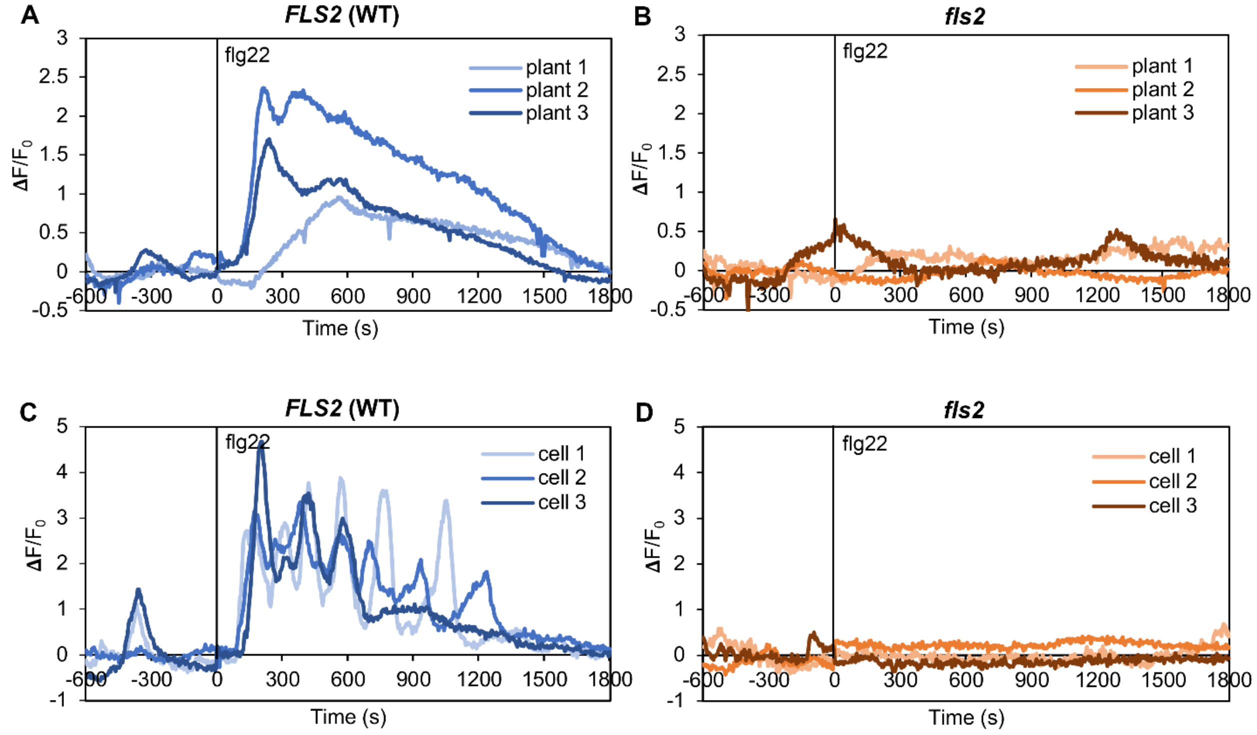


**Fig. S3. The flg22-induced cytosolic $Ca^{2+}$ oscillations are abolished in *fls2*.** (**A–D**) Fluorescence intensity ratio changes in cotyledon epidermal cells treated with 1 μM flg22 for the homozygous *fls2* mutant (B, D) or its wild-type sibling, *FLS2* (A, C) expressing R-GECO1. Fluorescence intensity profiles shown were calculated from the whole imaging field (A, B) or from individual cells (C, D). Data shown are from 3 cotyledons per genotype.

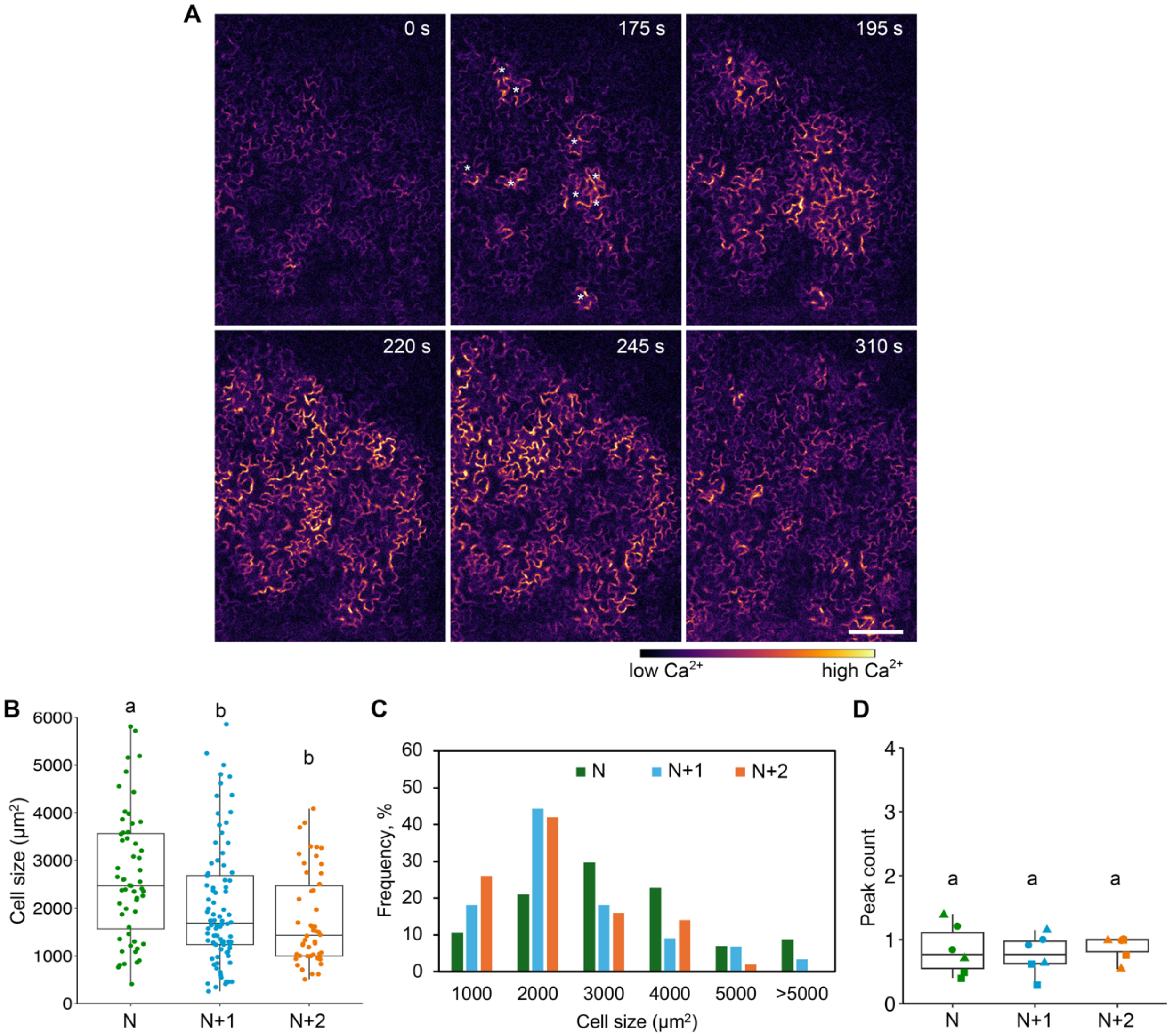


**Fig. S4. Initiator cells are randomly distributed in epidermal tissue and are larger than neighboring cells.** (**A**) Representative timelapse image series shows traveling wave propagation in cotyledon epidermal cells following global treatment with 1 µM flg22. See also Supplemental Movie S2. Initiator cells are marked with an asterisk. Bar = 100 µm. (**B, C**) Quantitative analysis of the area of initiator cells (N), N+1 and N+2 neighboring cells. Initiator cells were significantly larger than N+1 and N+2 cells. N = 57, 88, and 50 N, N+1 and N+2 cells from 8 cotyledons. One-way ANOVA and Turkey's HSD test, different letters indicate significant differences with $P < 0.01$. (**D**) Quantification of the number of $Ca^{2+}$ spikes during the 10 min run-in period for N, N+1 and N+2 cells. N = 6 cotyledons from 3 independent experiments. One-way ANOVA and Tukey's HSD test, $P > 0.05$.

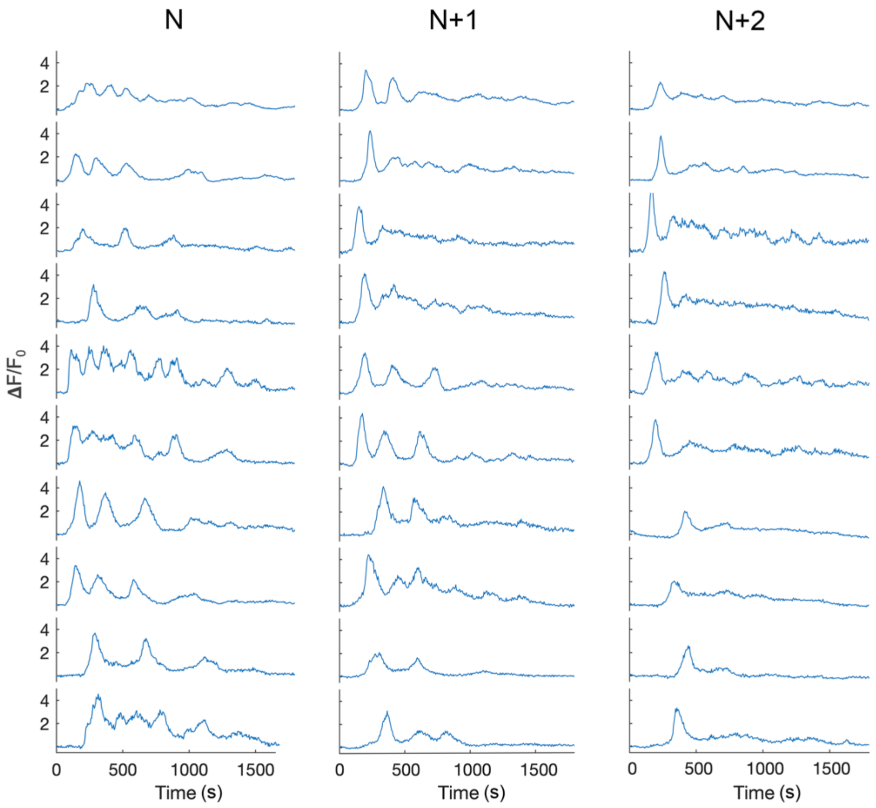


**Fig. S5. Representative traces of $Ca^{2+}$ intensity ratio ($\Delta F/F_0$) for initiator (N) and neighboring (N+1 and N+2) cells following global treatment with 1 µM flg22.** Representative traces are from the same dataset shown in Fig. 1. Time 0 represents when flg22 was added. Ten traces from 5 cotyledons are shown for each cell type.

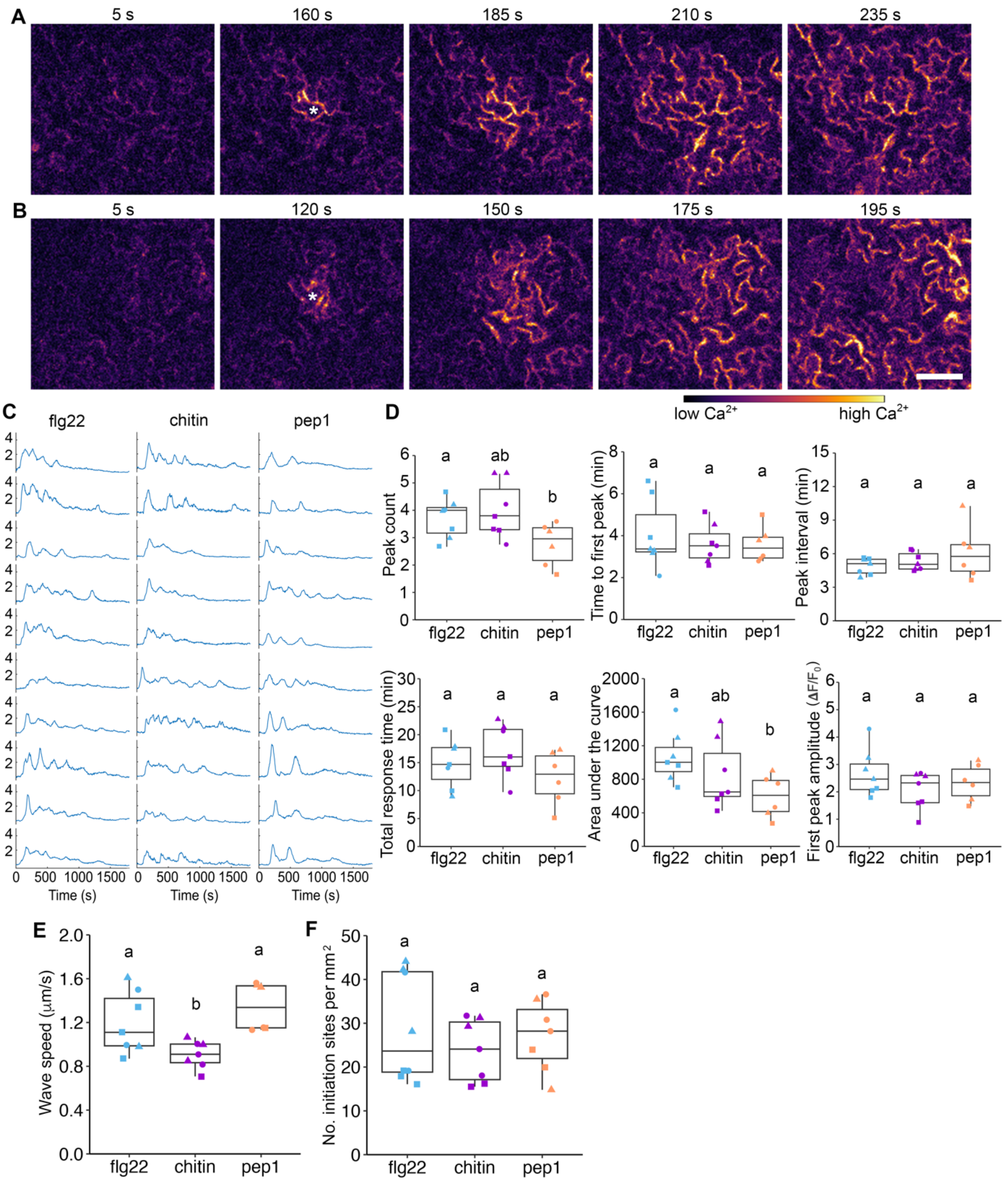


**Fig. S6. Calcium signatures and wave propagation patterns in cotyledon epidermal cells treated with various MAMPs or DAMPs.** (**A, B**) Representative timelapse image series showing traveling waves of $Ca^{2+}$ in cotyledon epidermal cells elicited by global treatment with 20 µg/mL chitin (A) or 100 nM Pep1 (B). See also Supplemental Movies S3 and S4. Time 0 represents when elicitors were added to the imaging chamber. Initiator cells are marked with an asterisk. Bar = 50 µm. (**C**) Representative intensity ratio traces ($\Delta F/F_0$) for initiator cells elicited by global treatment with 100 nM flg22, 20 µg/mL chitin or 100 nM Pep1. (**D**) Quantitative analysis of peak features for $Ca^{2+}$ traces from initiator cells. (**E**) Quantitative analysis of traveling wave speed in cotyledons treated with MAMPs or DAMPs. (**F**) Quantification of the density of initiation sites for traveling waves in cotyledon epidermis tissue. Each data point in the box plots represents an average value measured from 3–5 cells or 3–6 waves from a single cotyledon; data from 6–7 cotyledons from 3 independent experiments denoted by different shapes are presented in each box plot. One-way ANOVA and Tukey's HSD test, different letters indicate significant differences with $P < 0.05$.

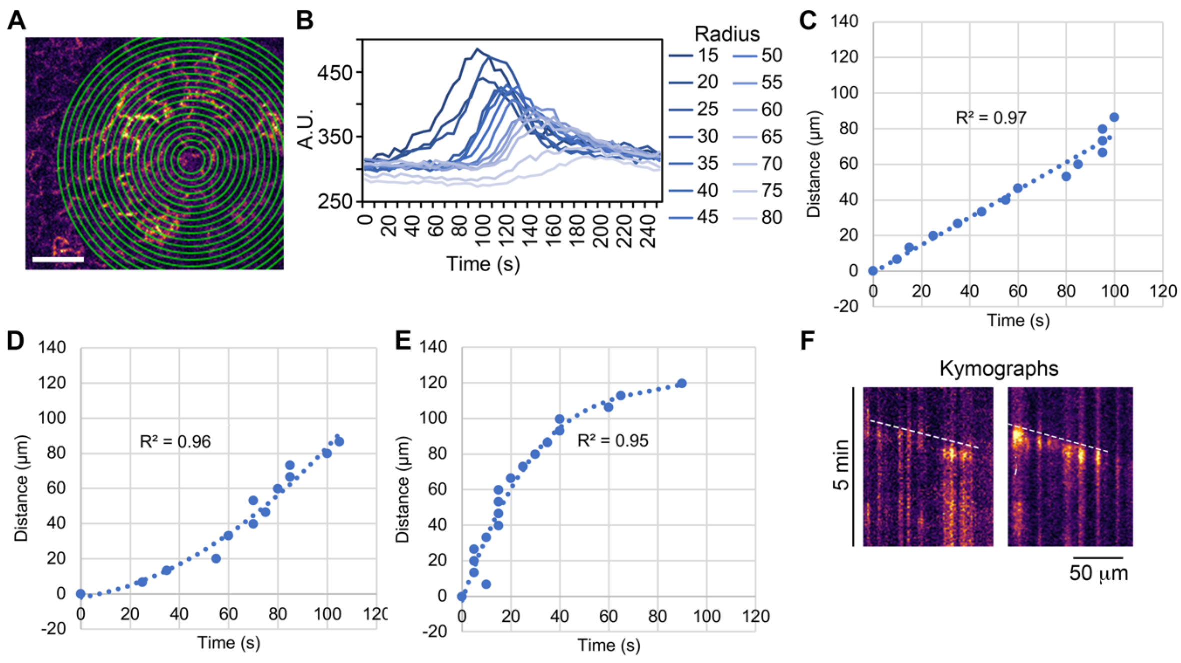


**Fig. S7. A method for analysis of the speed of intercellular $Ca^{2+}$ traveling waves in the cotyledon epidermis.** (**A**) A series of concentric circles, with the radius increased by 5 pixels at each step, was generated from the center of a traveling wave. Bar = 50 µm. (**B**) The mean fluorescence intensity for each circular ROI across the entire time series was extracted and a representative plot of the intensity profiles for all ROIs is shown in (B). The peak of each trace in (B) was identified based on its intensity and the corresponding value on the x-axis represents the time a wave travels a given distance (radius of the circle). (**C–E**) Representative plots of wave travel distance over time in cotyledons following global treatment with 1 µM flg22. Time was normalized so that t = 0 represents the initiation of the wave. Different patterns of scatter plots were fitted with a linear function in (C), a polynomial of degree 2 in (D), and a polynomial of degree 3 in (E). (**F**) Representative kymographs generated from the time series shown in Fig. 4A and 4B. A straight line could be fitted to the trajectory of the wave front (white dashed line), suggesting a constant speed.

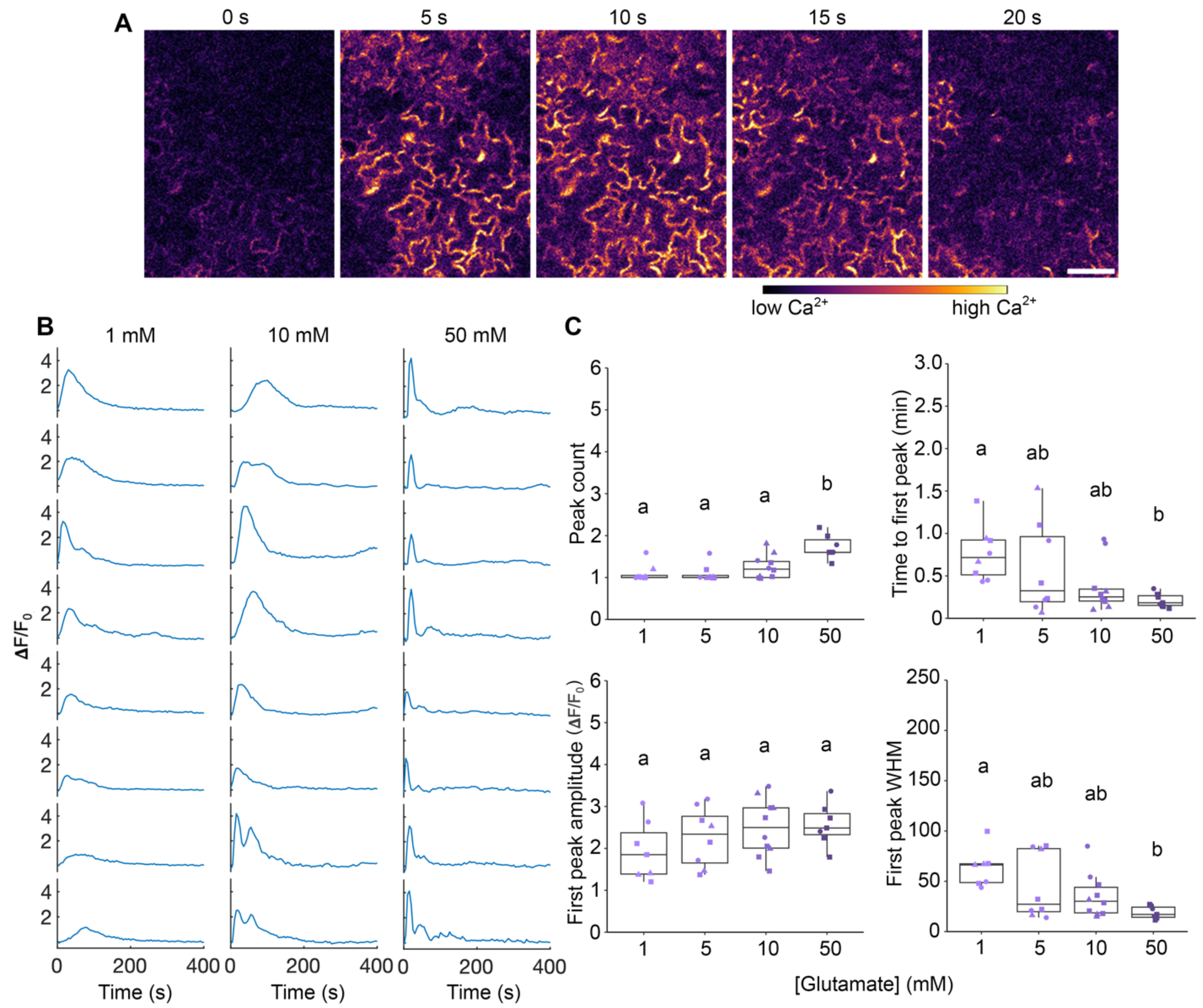


**Fig. S8. Single-cell $Ca^{2+}$ signatures in response to global glutamate treatment differ from those elicited by MAMPs.** (**A**) Representative timelapse image series shows $Ca^{2+}$ dynamics in cotyledon epidermal cells induced by global treatment with 10 mM glutamate. Time 0 represents when treatments were added to the imaging chamber. Bar = 50 µm. (**B**) Representative traces of $Ca^{2+}$ intensity ratio from single cells following global treatment with 1, 10, or 50 mM glutamate. (**C**) Quantitative analysis of peak features for $Ca^{2+}$ traces from cells treated with different doses of glutamate. WHM: width at half maximum. Each data point in the box plots represents an average value measured from 5 cells from a single cotyledon; data from 7–10 cotyledons from 3 independent experiments denoted by different shapes are presented in each box plot. One-way ANOVA and Tukey's HSD test, different letters indicate significant differences with $P < 0.05$.

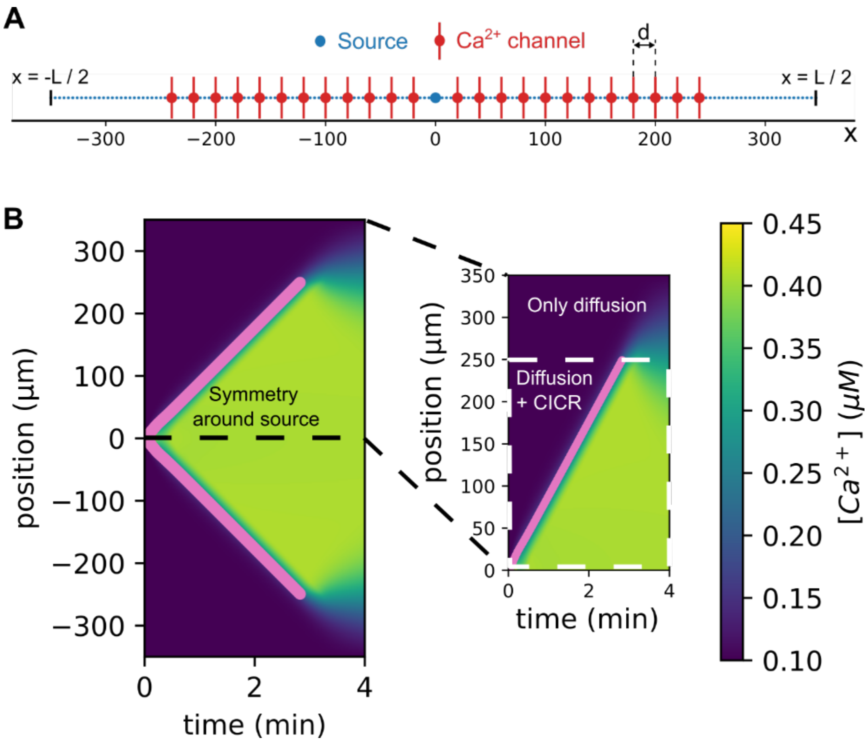


**Fig. S9. Mathematical modeling of 1D wave propagation through $Ca^{2+}$-induced $Ca^{2+}$ release.** (**A**) A representative 1D grid illustrating the model geometry, with a total length L = 700 µm. $Ca^{2+}$ channels are positioned between -250 µm and 250 µm, spaced by distance d, which is a model parameter. The center (x = 0) serves as the initiation site for $Ca^{2+}$ release. (**B**) A kymograph displaying changes in $Ca^{2+}$ concentration across various spatial points (Y-axis) over time (X-axis). Pink markers denote the time at which specific $Ca^{2+}$ channels are activated, representing the leading edge of the model-predicted calcium wave. Due to symmetry around the source site, only the half-domain from 0 to 350 µm is visualized in the panel on the right. The color scale within the kymograph represents $Ca^{2+}$ concentration, with values indicated by the bar on the right.

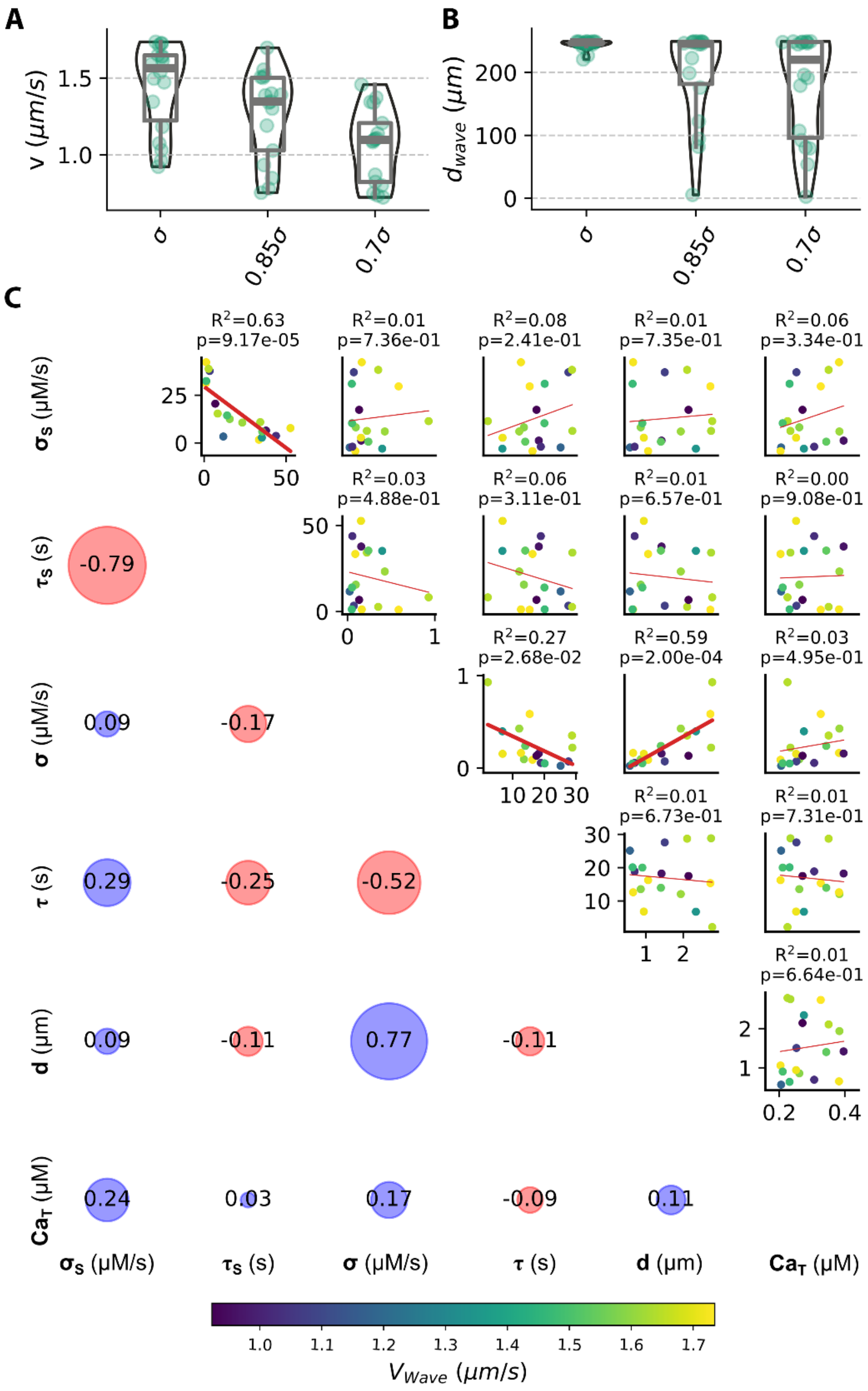


**Fig. S10. Local sensitivity and correlation analysis of model parameters.** (**A, B**) For each parameter set that produced wave speeds within the experimental range (Fig. 5A), the parameter $\sigma$ was reduced by 15% and 30% as shown on the x-axis. A box-and-whisker plot displays the distribution of wave speeds (A) and wave propagation distances (B) resulting from these reductions in $\sigma$. The box spans the interquartile range (IQR), with whiskers extending to 1.5 times the IQR, and the horizontal line within the box indicates the median. (**C**) Correlation analysis of the parameter values for the best-fitting parameters from Fig. 5A. The upper diagonal shows scatter plots for each parameter pair, color-coded by wave speed. The line of best fit is overlaid in red color, with the $R^2$ value indicated above each plot. The lower diagonal shows the Pearson correlation coefficients, represented by circles. The size and color of each circle correspond to the strength and direction of the correlation: red for negative and blue for positive correlations.

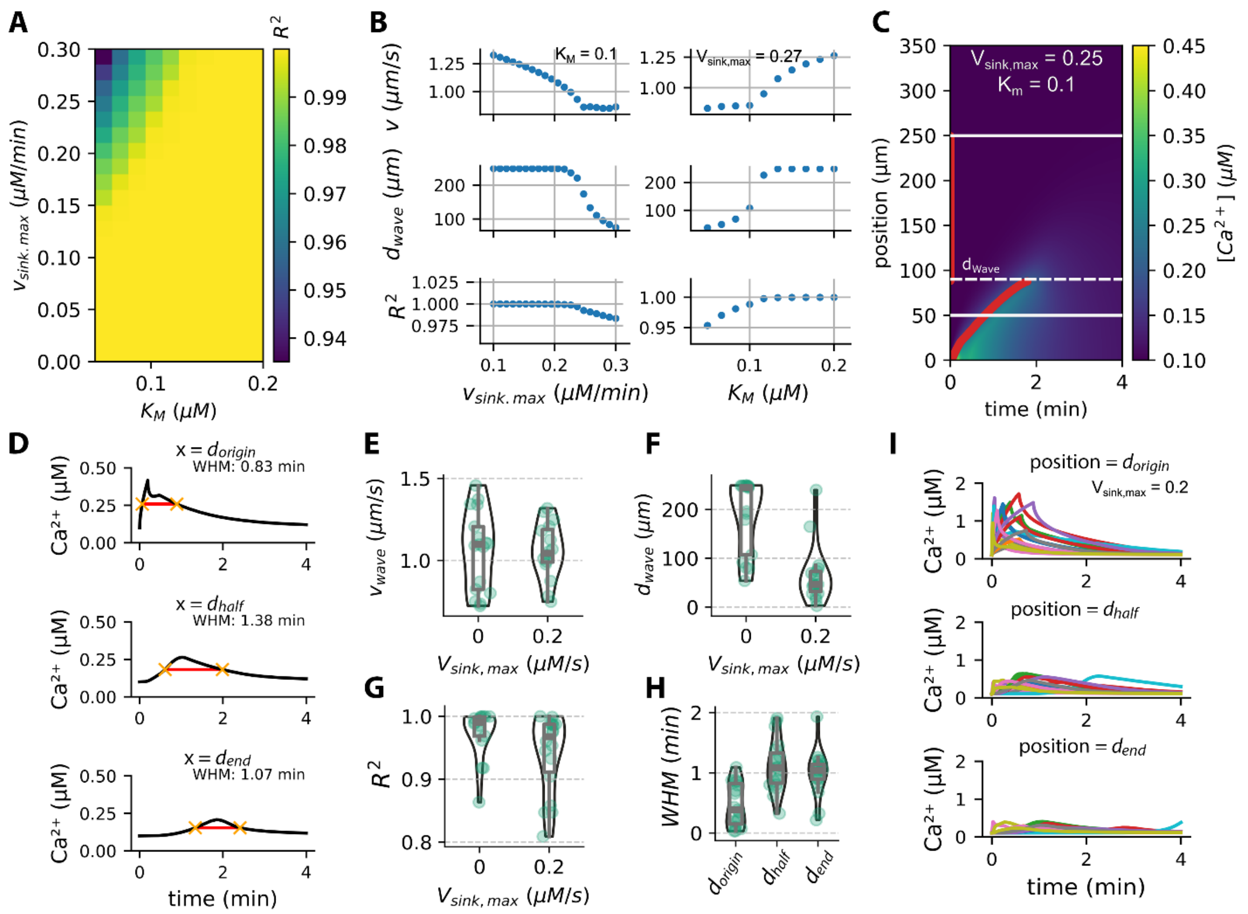


**Fig. S11. Extended analysis for mathematical model with a sink.** (**A**) Parameters $K_M$ and $V_{sink,max}$ were varied within specified ranges. For each parameter combination, we calculated the $R^2$ values represented as a heatmap by fitting a line to the distances of CICR channels from the initiator site, plotted against their activation times. Base parameter values of the CICR model used in these studies were taken from a representative calibrated parameter set and are $\sigma_S$ = 3.34, $\tau_S$ = 11.71 s, $\sigma$ = 0,035 μM/s, $\tau$ = 25.08 s, $d$ = 0.57 μm, $Ca_T$ = 0.21 μM and $Ca_b$ = 0.1 μM (**B**) For the base parameter set $K_M$ fixed as 0.1 μM, we varied the parameter $V_{sink,max}$ and visualized the resulting changes in wave speed ($v$), distance travelled by wave before attenuation ($d_{wave}$) and $R^2$ of the linear fit with increasing $V_{sink,max}$. A similar analysis was performed by varying parameter $K_M$ and keeping parameter $V_{sink,max}$ fixed at 0.27 μM/min. (**C**) Kymograph showing $Ca^{2+}$ concentration in the one-dimensional simulation domain over time (x-axis), with distance along the domain on the y-axis. A sink was introduced in the fire-diffuse-fire model with parameters $V_{sink,max}$ = 0.25 μM/min, $K_M$ = 0.11 μM, and n = 2. (**D**) $Ca^{2+}$ concentration is plotted over time from positions of $Ca^{2+}$ channels near the origin (top), half (middle), and endpoint (bottom) of the total wave propagation distance. A horizontal red line indicates the half-maximum width (WHM) within each plot. (**E–G**) For each parameter set producing wave speeds within the experimental range (Fig. 5A), the parameter $V_{sink,max}$ was added to the model and set to a value of 0.2 μM/min as shown on the x-axis. A box-and-whisker plot displays the distribution of (E) wave speeds ($v$), (F) wave propagation distances ($d_{wave}$), and (G) $R^2$ values resulting from the inclusion of a sink in the model. The box spans the interquartile range (IQR), with whiskers extending to 1.5 times the IQR, and the horizontal line within the box indicates the median. (**H**) For each parameter set where a sink was added, we analyzed the half-maximal width for $Ca^{2+}$ signals at sites near the initiation site or source (origin), at the midpoint of the wave's propagation (half), and at the wave's endpoint (end). A combination of box and violin plots visualizes the distributions of half-maximal widths at these locations. (**I**) Representative $Ca^{2+}$ signals at three different sites shown in (H). Each color shows a fit for a particular parameter value. Location of site with respect to the distance traveled by the wave is indicated on top of each panel.

**Table S1. List of CICR model parameters**

| Parameter | Definition | *Xenopus* eggs (Dawson et al., 1999) | *Xenopus* oocytes (Dawson et al., 1999) | *Arabidopsis* roots (Evans et al., 2016) | 1D CICR model parameter bounds |
|---|---|---|---|---|---|
| $\sigma_S$ | Source/Initiator site flux | - | - | - | [1.6e-05, 52.70] μM/s |
| $\tau_S$ | Source activation time | - | - | - | [0.06–60] s |
| $D$ | Diffusion coefficient | 50 $\mu m^2/s$ | 25 $\mu m^2/s$ | 20 $\mu m^2/s$ | 20 $\mu m^2/s$ |
| $\sigma$ | Activation strength of $Ca^{2+}$ channel | 0.17 μM/s | 47.58 μM/s | - | [1.6e-05, 52.70] μM/s |
| $\tau$ | Activation time of $Ca^{2+}$ channel | 9 s | 0.05 | - | [0.06–60] s |
| $d$ | Distance between two $Ca^{2+}$ channels | 3 μm | 3.9 μm | 1 μm | [0.5–3] μm |
| $V_{sink,max}$ | Maximum flux of $Ca^{2+}$ through sink | - | - | - | [0-5e–3] μM/s |
| $K_m$ | $Ca^{2+}$ concentration for half saturation of sink type proteins | - | - | - | [0.05–0.3] μM |
| $n$ | Number of $Ca^{2+}$ ions needed for sink activation | - | - | - | 2 |
| $Ca_b$ | Basal $Ca^{2+}$ concentration | - | 0.05 μM | 0.3 μM | 0.1 μM |
| $Ca_T$ | Threshold $Ca^{2+}$ concentration for activation of $Ca^{2+}$ channel | - | 0.25 μM | 0.4 μM | [0.2–0.4] μM |
| $v_{wave}$ | Wave speed | 5.2 μm/s | 20 μm/s | 396 ± 28 μm/s | |

### Supplemental Movie Legends

**Movie S1. Cytosolic $Ca^{2+}$ dynamics in cotyledon epidermal cells following global treatment with flg22.** Same as Figure 1A. Time 0 represents when 1 µM flg22 was added to the imaging chamber. Video playback rate = 20 frames per second (fps). Total elapsed time = 2400 s. Bar = 50 µm.

**Movie S2. Local traveling waves of $Ca^{2+}$ induced by flg22 are initiated from a small subset of cells randomly distributed in epidermal tissue.** Same as Figure S4A. Time 0 represents when 1 µM flg22 was added to the imaging chamber. Video playback rate = 20 fps. Total elapsed time = 2400 s. Bar = 50 µm.

**Movie S3. Cytosolic $Ca^{2+}$ dynamics in cotyledon epidermal cells following global treatment with chitin. Same as Figure S6A.** Time 0 represents when 20 µg/mL chitin was added to the imaging chamber. Video playback rate = 20 fps. Total elapsed time = 2400 s. Bar = 50 µm.

**Movie S4. Cytosolic $Ca^{2+}$ dynamics in cotyledon epidermal cells following global treatment with Pep1.** Same as Figure S6B. Time 0 represents when 100 nM Pep1 was added to the imaging chamber. Video playback rate = 20 fps. Total elapsed time = 2400 s. Bar = 50 µm.

**Movie S5. Wounding of a single epidermal cell with laser irradiation induces a $Ca^{2+}$ traveling wave. Same as Figure 3A.** Time 0 represents when the laser was applied to the epidermal cell. Video playback rate = 20 fps. Total elapsed time = 240 s. Bar = 50 µm.

**Movie S6. Calcium traveling waves in cotyledon epidermal cells elicited by 10 nM flg22. Same as Figure 4B.** Time 0 represents when 10 nM flg22 was added to the imaging chamber. Video playback rate = 20 fps. Total elapsed time = 1800 s. Bar = 50 µm.
